# Revealing how variations in antibody repertoires correlate with vaccine responses

**DOI:** 10.1101/2021.08.06.454618

**Authors:** Yana Safonova, Sung Bong Shin, Luke Kramer, James Reecy, Corey T. Watson, Timothy P.L. Smith, Pavel A. Pevzner

## Abstract

An important challenge in vaccine development is to figure out why a vaccine succeeds in some individuals and fails in others. Although antibody repertoires hold a key to answering this question, there have been very few personalized immunogenomics studies so far aimed at revealing how variations in immunoglobulin genes affect a vaccine response. We conducted an immunosequencing study of 204 calves vaccinated against bovine respiratory disease (BRD) with the goal to reveal variations in immunoglobulin genes and somatic hypermutations that impact the efficacy of vaccine response. Our study represents the largest longitudinal personalized immunogenomics study reported to date across all species, including humans. To analyze the generated dataset, we developed an algorithm for identifying variations of the immunoglobulin genes (as well as frequent somatic hypermutations) that affect various features of the antibody repertoire and titers of neutralizing antibodies. In contrast to relatively short human antibodies, cattle have a large fraction of ultralong antibodies that have opened new therapeutic opportunities. Our study revealed that ultralong antibodies are a key component of the immune response against the costliest disease of beef cattle in North America. The detected variants of the cattle immunoglobulin genes, which are implicated in the success/failure of the BRD vaccine, have the potential to direct the selection of individual cattle for ongoing breeding programs.

## Introduction

### The challenge of identifying variations in immunoglobulin genes that affect vaccine response

Although vaccination is a primary tool to control the spread of viral and bacterial diseases, the success of vaccines at the population level does not always translate to protection at the individual level. Figuring out why a vaccine fails in some individuals is important both during the vaccine development stage (to inform changes in the vaccination protocols) and the vaccine administration stage (to identify a sub-population with a poor vaccine response). A promising approach to understanding why a vaccine succeeds in some individuals and fails in others is to analyze the germline variations in the immunoglobulin (IG) loci of individuals with successful/failing antibody responses to the vaccine.

Antibodies are not encoded in the germline genome but rather result from somatic genomic recombinations called *VDJ recombinations* (Tonegawa, 1983). This process affects an IG locus containing the families of the *variable* (*V*), *diversity* (*D*), and *joining* (*J*) genes (referred to as *IG genes*) by selecting one V, one D gene, and one J gene, and concatenating them together to generate one of the antibody chains. Further diversity of antibodies is generated by the class-switch recombination and somatic hypermutations (SHMs) (Dudley et al. 2005). There are three types of IG loci in mammalian species (including cows): heavy chain, kappa light chain, or lambda light chain. In this work, we focus on the heavy chain (IGH) locus only.

The *expression quantitative trait loci* (*eQTL*) analysis links variation in gene expression to genotypes. Although eQTL analysis has greatly contributed to the dissection of the genetic basis of disease and vaccine response (Franco et al., 2013, Bhalala et al., 2018), the IG loci remain virtually untouched by eQTL studies (Watson et al., 2017). eQTL studies usually start from generating an *n*×*m genotype matrix* that contains information about each of *m* markers (e.g., SNPs) in each of *n* individuals and an *n*×*k phenotype* (*expression) matrix* that contains information about expression levels of each of *k* genes in each of *n* individuals. Generating analogs of the genotype and phenotype matrices in immunogenomics studies is a more complex task than in traditional eQTL studies.

First, while the set of genes in eQTL studies is fixed and shared by all individuals, the antibody repertoire is composed of a virtually unlimited set of proteins, and there are typically few antibodies shared between any two individuals. Thus, given an antibody repertoire represented as a *repertoire sequencing* (*Rep-seq*) dataset, it is not clear how to define the phenotype matrix. One possibility is to consider each germline gene in the IG locus (e.g., a V gene, a D gene, or a J gene) and to define the *usage* of this gene as the fraction of antibodies that originated from this gene among all antibodies in the Rep-Seq dataset. Our goal is to identify *usage QTLs* (*IgQTLs*) that link variation in usage to a genotype.

Second, eQTL studies are usually based on RNA-seq and Whole Genome Sequencing (WGS) data, while immunosequencing studies generate Rep-seq data about antibodies. Thus, the genotype matrix in immunosequencing studies has to be inferred from Rep-seq data alone since the WGS data is typically not available. Although inference of alleles of V, D, and J genes from Rep-seq data is a well-studied problem (Gadala-Maria et al., 2015; Corcoran et al., 2016; Gadala-Maria et al., 2019; Safonova and Pevzner, 2019; Bhardwaj et al., 2020), the existing allele inference tools, primarily developed for naïve repertoires, often fail in the case of more complex antigen-stimulated repertoires that represent the primary goal of IgQTL studies. Also, in contrast to allele inference tools that attempt to infer SNPs and ignore frequent somatic hypermutations, IgQTL studies should account for both SNPs and frequent SHMs since they may play equally important roles in vaccine responses.

### Bovine Respiratory Disease

We conducted a personalized immunogenomics study of 204 calves to analyze the efficacy of the bovine respiratory disease (BRD) vaccine, the largest time-series immunosequencing dataset generated so far across all species, including human. Since cattle production accounted for $67 billion in 2018 in the United States (Economic Research Service USDA, 2021), maintaining cattle health is an important direction of agricultural studies. The BRD is the costliest disease of beef cattle in North America (Taylor et al., 2010). Although vaccination reduces the risk of BRD, losses from BRD remain substantial and individuals respond very differently to the BRD vaccine (Kramer et al., 2017). In order to understand links between variants in cattle IG genes and antibody responses to the vaccine, we generated four Rep-Seq datasets (taken before and after the BRD vaccination) for each of 204 calves.

### Ultralong cattle antibodies

The evolution of the cattle IGH locus has resulted in a loss of many functional V genes, thus reducing the diversity of the cattle antibody repertoire (Haakenson et al., 2018). To compensate for the reduced VDJ recombination diversity, cattle have developed antibodies with ultralong CDR3s that have a unique mechanism of *structural diversification* (Dong et al., 2019). We refer to antibodies with CDR3s longer than 50 amino acids as *ultralong antibodies* (for comparison, the average length of human CDR3s is only 15 aa). Although non-conventional recombination processes (such as D-D fusions and V gene replacement) can generate ultralong human antibodies (including, broadly neutralizing human antibodies against HIV-1, Yu and Guan, 2014), they are rare in human antibody repertoires, typically accounting for less than 1% of all antibodies (Safonova and Pevzner, 2019). In contrast, ultralong antibodies account for ∼10% of cattle antibodies (Wang et al., 2013).

The vast majority of ultralong cattle antibodies are generated by the VDJ recombination of the same V, D, and J genes: IGHV1-7, IGHD8-2, and IGHJ2-4 (Wang et al., 2013). An unusually long IGHD8-2 gene (148 nt) contributes to all ultralong CDR3s and enables their structural diversification. This gene encodes four cysteines that form disulfide bonds and turn the CDR3 loop into a complex protein structure called a *knob* (Wang et al., 2013). IGHD8-2 also contains many codons that differ from the cysteine codons by a single nucleotide. SHMs often result in new cysteines that form new disulfide bonds, thus extending the diversity of the knob structures.

Ultralong cattle antibodies open new therapeutic opportunities (Muyldermans and Smider, 2016) and may even neutralize various strains of HIV (Sok et al., 2017). Human antibodies target the HIV envelope glycoprotein (*Env*) presented on the virus surface. However, highly mutated *Env* proteins are often covered by glycans, making them a hard-to-reach target for human antibodies. Ultralong cattle antibodies penetrate glycans and directly target the conservative sites that are unreachable for human antibodies since they are buried inside the *Env* protein. Although ultralong cattle antibodies have great pharmaceutical potential in terms of targeting unusual antigens (Burke et al., 2020), their role in the native immune response remains unclear.

### The challenge of finding IgQTLs

Previous immunosequencing studies have succeeded in linking variants in the human IGH locus to disease and vaccination efficacy (Thomson et al., 2008; Lingwood et al, 2012; Avnir et al., 2016; Parks et al., 2017, Lee et al., 2021, Mikocziova et al., 2021) but have not yet resulted in a software tool addressing the above-mentioned complications in finding IgQTLs. We thus developed the IgQTL tool for detecting both variants of germline V genes and their frequent SHMs for downstream analysis of usage QTLs in the V genes. Using IgQTL, we inferred genotypes associated with the fraction of ultralong antibodies in a repertoire, and the fraction of antibodies specific to BRD antigens. Our findings indicate that ultralong antibodies play an important role in the bovine antibody response against BRD antigens and suggest that it is important to add immunosequencing to the existing genomics-based breeding efforts in the cattle industry.

## Results

### Rep-Seq datasets

We analyzed Rep-seq datasets for 204 purebred American Angus calves vaccinated against BRD. Each animal was initially vaccinated at day 0 and then given a booster vaccination three weeks later (Figure 1A). For each animal, four IgG samples were collected: three weeks prior to vaccination (referred to as “–3”), at the moment right after the first vaccination (referred to as “0”), three weeks post-vaccination (referred to as “+3”), and six weeks post-vaccination (referred to as “+6”). Serum from each of the sequenced animals was also assayed for BRD-specific antibody titers at the same time points (Kramer et al., 2017) to quantify pre-existing immunity (e.g., resulting from maternal antibodies that are passed through milk from the mother to the child) and as a measure of vaccine success.

**Figure 1.**
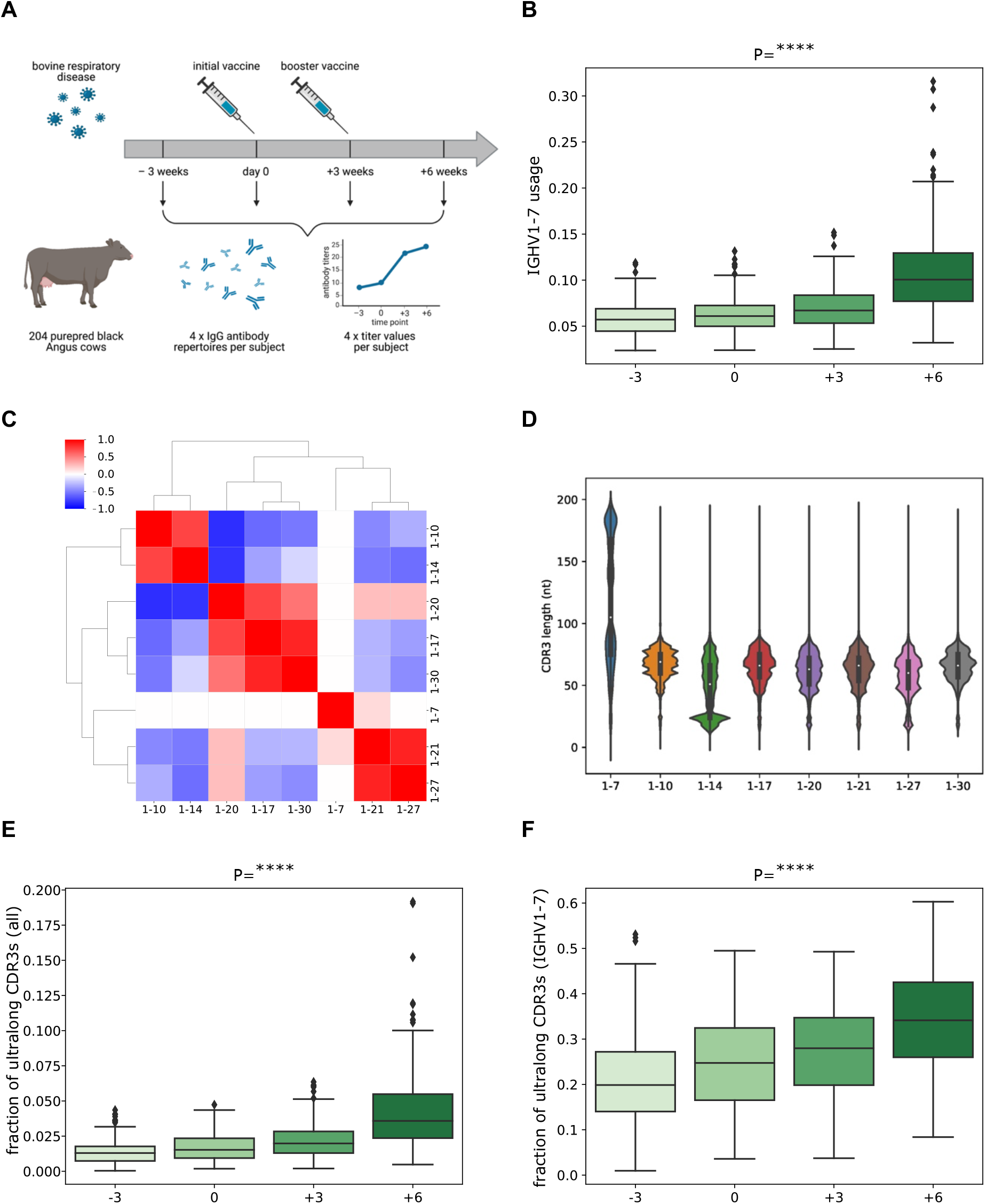
Overview of study design and characteristics of antibody repertoires. (*A*) The panel shows an overview of the study design. 204 calves were vaccinated against BRD, and their expressed antibody repertoires were sequenced at four time points pre- and post-vaccination. Serum from each of the sequenced animals was also assayed for BRD-specific antibody titers at the same time points. (*B*) The distribution of IGHV1-7 usage at four time points. Here and further, each box shows the quartiles of the distribution. The whiskers show the rest of the distribution, except for outliers found using a function of the inter-quartile range implemented by the Seaborn package in Python. P-values have the following notation: ns≥0.05, *<0.05, **<0.01, ***<0.001, ****<0.0001. (*C*) The matrix shows the Pearson correlations between gene usages computed across all high-usage V genes at time point “–3”. Correlation values vary from –1 (blue) to 1 (red). Statistically insignificant correlations (P≥0.05) are shown as white cells. (*D*) The histogram of the distributions of the CDR3 lengths for eight highly-used cattle V genes. The histogram is computed for individual 14007. (*E*) The distribution of the fraction of ultralong CDR3s in all CDR3s at four time points. (*F*) The distribution of the fraction of ultralong CDR3s in CDR3s derived from IGHV1-7 at four time points.

### High-usage cattle V genes

Each Rep-Seq dataset was aligned to 13 cattle V genes from IMGT, the international ImMunoGeneTics information system (Lefranc et al., 2009) (see Methods). For each V gene, we computed its usage in an individual as the fraction of sequences with distinct CDR3s aligned to this V gene in the corresponding Rep-Seq dataset. We defined the *average usage* of a V gene as its average usage across all individuals and all time points. Only 8 out of 13 V genes had an average usage exceeding 0.01: IGHV1-7, IGHV1-10, IGHV1-14, IGHV1-17, IGHV1-20, IGHV1-21, IGHV1-27, and IGHV1-30. We refer to them as *high-usage* V genes and limit further analysis to these genes only.

### IGHV1-7 is the only V gene with a statistically significant increase in usage after vaccination

We first assessed differences in usage between pre- and post-vaccination for all V genes. We found that only IGHV1-7 has significantly higher usage after the vaccination (Table S1). Figure 1B shows that the usage of the IGHV1-7 gene is significantly increased at time points “+3” and “+6” (P-value=8.17×10^−115^), suggesting that vaccination triggered the production of antibodies derived from IGHV1-7. Henceforth, we used the linear mixed effect model to estimate P-values (referred to simply as “P”) for repeated measures (representing four time points), unless a different method is specified.

Analysis of Rep-seq datasets collected at the time point “–3” revealed that each V gene but IGHV1-7 has positive or negative correlation of usages with other IGHV genes. Figure 1C shows that V genes form three groups based on their usage, consisting of seven positively correlated genes: G1=(IGHV1-10, IGHV1-14), G2=(IGHV1-17, IGHV1-20, IGHV1-30), and G3=(IGHV1-21, IGHV1-27). IGHV1-7 is the only gene with an independent usage profile. These correlations are consistent across time points “0”, “+3”, and “+6”. We conjecture that this is explained by an association of IGHV1-7 with ultralong CDR3s and their special role in the adaptive immune response. Figure 1C illustrates that the usages in groups G1, G2, and G3 anticorrelate.

### BRD vaccination triggers the increased production of ultralong antibodies

Figure 1D shows that for all high-usage cattle V genes, except for IGHV1-7, the mean CDR3 length does not exceed 75 nt. It also illustrates that, unlike other V genes, IGHV1-7 has a bimodal distribution of CDR3 lengths, and the second mode is represented by ultralong CDR3s. On average, 21% of sequences derived from IGHV1-7 have ultralong CDR3s at time point “–3” and 99% of ultralong CDR3s across all individuals are derived from IGHV1-7. The latter observation agrees with findings reported by Walther et al., 2013, Wang et al., 2013, Deiss et al., 2017, and Dong et al., 2019.

Figure 1E illustrates that both initial and booster vaccinations significantly increase the fraction of ultralong CDR3s in all CDR3s (across all V genes) at time points “+3” and “+6” (P=3.61×10^−107^). Figure 1F shows that the fraction of ultralong CDR3s in all CDR3s derived from IGHV1-7 also increases after vaccinations (P=1.19×10^−119^). These observations suggest that the vaccination triggers the production of antibodies derived from IGHV1-7.

### Antibody titers correlate with fractions of ultralong CDR3s

Some calves have pre-existing immunity because they either were previously exposed to the BRD-causing virus or have maternal antibodies specific to BRD. This pre-existing immunity (as well as cross-reactivity of antibodies) may affect titers at the initial time point “–3”. Downey et al., 2013 demonstrated that the decay rate of maternal antibodies is rather low and that there is a threshold effect: the calves do not respond to the vaccine if the level of maternal antibodies exceeds a threshold and only respond when this level drops. Also, the impact of calf age on antibody response (titers) to BRD was previously shown to be insignificant (Kramer et al., 2017).

Figure 2A shows that, on average, the booster vaccination increased neutralizing antibody titers. Figure 2B shows the Pearson correlation *r* between antibody titers at four time points and illustrates that they correlate at points “–3” and “0” (*r*=0.78, P=5.83×10^−43^), “–3” and “+3” (*r*=0.43, P=1.53×10^−10^), and “0” and “+3” (*r*=0.43, P=2.29×10^−10^). In contrast, antibody titers from time points “–3” and “0” anticorrelate with final titers at the time point “+6” (*r*=–0.42, P=3.34×10^−10^ and *r*=–0.4, P=1.78×10^−9^, respectively). This suggests that pre-existing immunity to BRD antigens may be suboptimal, preventing development of a successful immune response to the BRD vaccine. Impacts of suboptimal antibody responses caused by pre-existing immunity were reviewed by Zimmermann and Curtis, 2019 (for various antigens) and Iwasaki and Yang, 2020 (for SARS-CoV-2) and discussed in Supplemental Note “The relations between pre- and post-vaccination immunity to the BRD vaccine in calves”.

**Figure 2.**
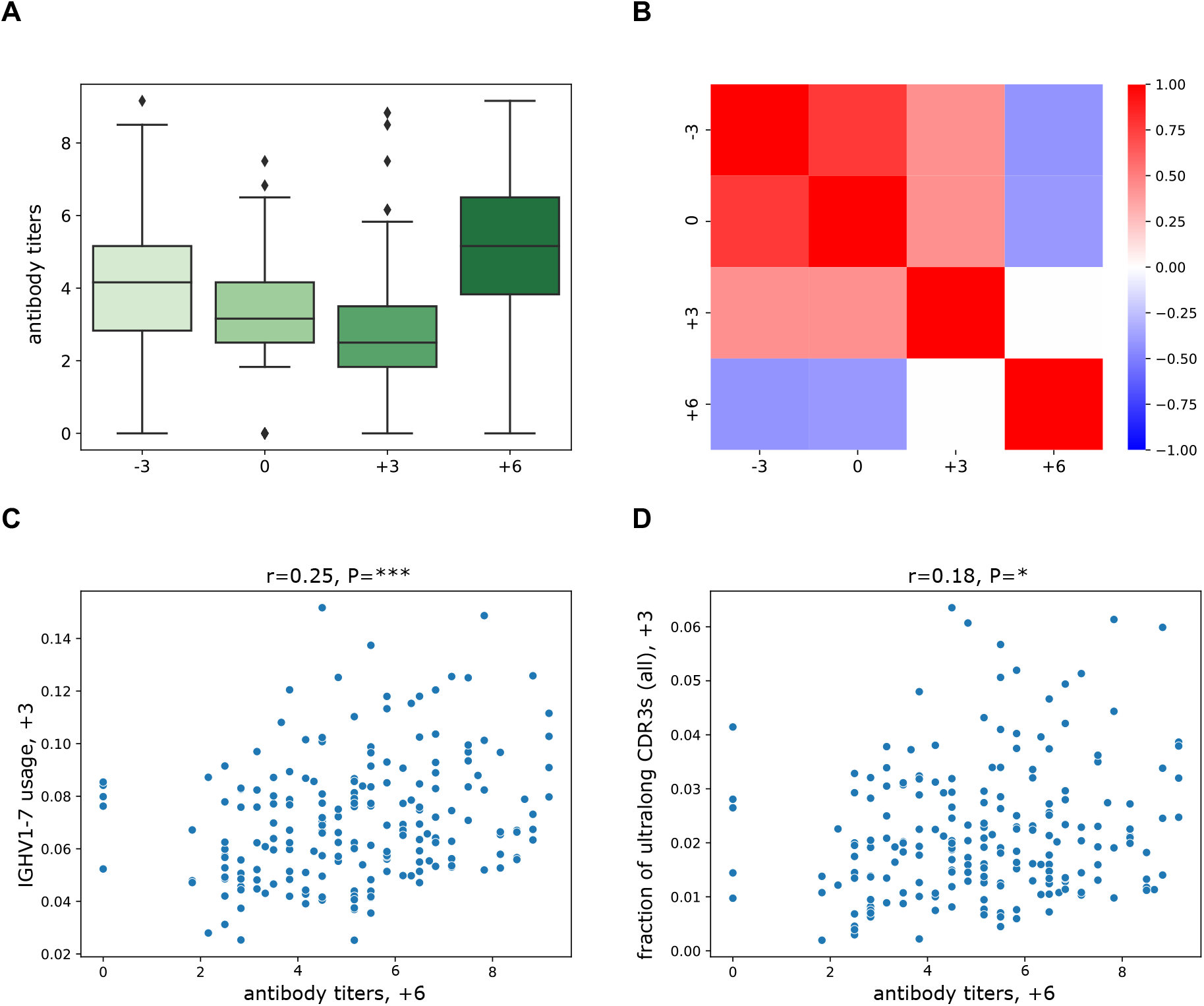
Antibody titers statistics. (*A*) The distribution of antibody titers at four time points. Titers at time point “-3” show the number of BRD-specific antibodies without any antigen stimulation. Titers at time point “0” represent immediate memory responses triggered by the vaccine. Titers at time points “+3” and “+6” reflect antibodies produced as a result of the vaccination. Details of the titer analysis are described in Kramer et al., 2017. (*B*) The matrix shows the Pearson correlations between antibody titers at four time points. Correlation values vary from –1 (blue) to +1 (red). Statistically insignificant correlations (P≥0.05) are shown as white cells. (*C*) Antibody titers at time point “+6” vs usages of IGHV1-7 at time point “+3”. (*D*) Antibody titers at time point “+6” vs fractions of ultralong CDR3s in all CDR3s at time point “+3”. The Pearson correlations (*r*) and P-values (P) are shown at the top of panels (*C*) and (*D*).

We have not found any statistically significant correlations between the titers and the usages of all high-usage V genes, except for IGHV1-7. Both the usage of IGHV1-7 (Figure 2C) and the fraction of ultralong CDR3s (Figure 2D) at the time point “+3” correlate (albeit weakly) with final titers at the time point “+6” (*r*=0.25, P=0.0004, and *r*=0.18, P=0.0125, respectively). These observations support our hypothesis that antibodies with ultralong CDR3s play an important role in recognizing the vaccine antigens.

### The IgQTL pipeline

To reveal the associations between germline/somatic variants and features of cattle antibody repertoires (gene usages, antibody titers, and fractions of ultralong antibodies), we developed the IgQTL tool. IgQTL takes Rep-Seq reads and antibody titers (if available) as an input and consists of the following steps (Figure 3A):

- Generating the phenotype matrix containing information about gene usages, fractions of ultralong CDR3s, antibody titers, etc.
- Finding germline and somatic variations (GSVs).
- Generating and clustering a genotype matrix to reveal subjects with common genotypes.
- Finding statistically significant genotype-phenotype associations.
- Identifying the most consequential GSVs with respect to phenotypes.

**Figure 3.**
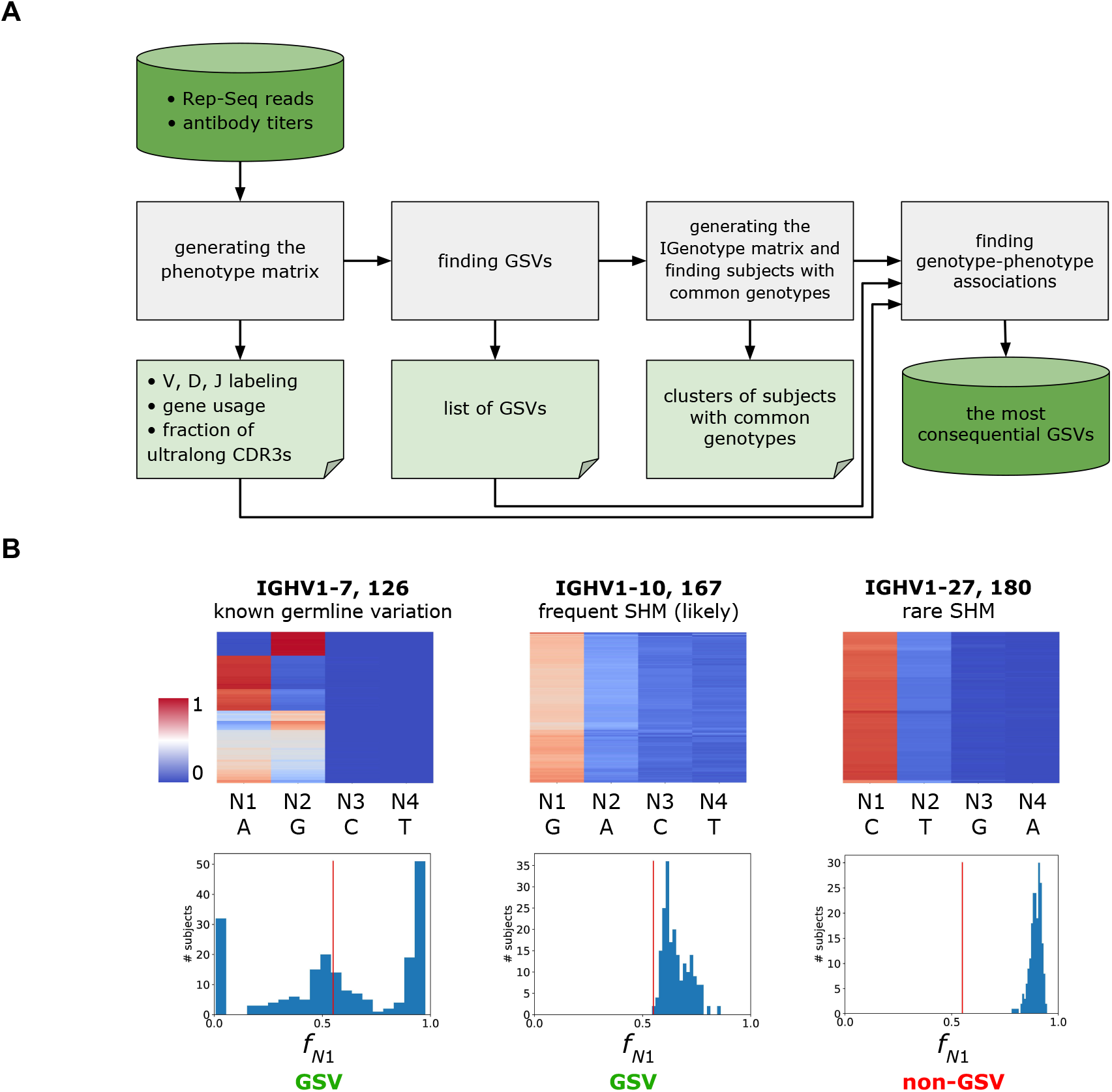
Overview of IgQTL method. (*A*) The panel illustrates IgQTL pipeline. Grey rectangles show various steps of the IgQTL pipeline. The input (Rep-Seq reads and antibody titers) and the final output (the most consequential GSVs) are shown in bright green. Intermediate output is shown in light green. (*B*) The panel illustrates the procedure for finding GSVs (126, IGHV1-7, A/G) and (167, IGHV1-10, G/A), as well as non-GSV (180, IGHV1-27, C/T). Heatmaps in the upper row show the fractions of the nucleotides across all subjects varying from 0 (blue) to 1 (red). Columns are arranged according to the sum of fractions across all subjects. *N*1 and *N*2 correspond to the first and the second columns, respectively. Histograms in the lower row show distributions of fractions *f*_*N*1_ across 204 animals. The red vertical line in each histogram corresponds to *freq*=0.55.

Below we applied IgQTL to reveal the genotype-phenotype associations in the cattle immunosequencing study.

### Generating the phenotype matrix

The phenotype matrix is defined as a matrix with 204 rows, where each column represents either the usage of one of V genes, or the fraction of ultralong antibodies, or an antibody titer. In the case of the cattle immunosequencing dataset, IgQTL forms a 204×14 phenotype matrix that represents usages of all high-usage V genes (the first eight columns), the fraction of ultralong antibodies among all antibodies in the repertoire (9^th^ column), the fraction of ultralong antibodies derived from IGHV1-7 among all antibodies derived from IGHV1-7 (10^th^ column), and the antibody titers (the last four columns).

### Finding germline variations and frequent SHMs in the V genes

Previous studies have revealed associations between germline variations in V genes and variation in their usages and antibody titers (Thomson et al., 2008; Lingwood et al, 2012; Avnir et al., 2016; Parks et al., 2017, Lee et al., 2021, Mikocziova et al., 2021). However, these studies did not consider the impact of frequent SHMs that, similarly to germline variations, may be associated with variation in gene usage. Since the Rep-Seq data, obtained from IgG antibodies, represent mature antibody responses, below we analyzed the impact of frequent SHMs on antibody repertoires.

To capture both germline variations and frequent somatic hypermutations (further referred to as *germline or somatic variations* or *GSVs*) for each subject, we generated a *combined* dataset from all four time points by collapsing identical sequences and analyzed sequences aligned to the same V gene. Given a position in a germline V gene, we analyzed all reads aligning to this gene in a single combined dataset and computed a vector (*f*_*A*_, *f*_*C*_, *f*_*G*_, *f*_*T*_), where *f*_*N*_ is the fraction of reads that have the nucleotide *N* aligned at this position. We collected such vectors from all subjects and define *N*1 and *N*2 as nucleotides with the highest and the second-highest total fractions.

For most positions, *f*_*N*1_ is close to 1, indicating that these positions do not exhibit variations and frequent SHMs. We were interested in positions where *f*_*N*1_ falls below a *frequency threshold freq* (the default value *freq*=0.55) as such positions likely reflect one of the following situations:

- if a subject is homozygous by *N*2 (i.e., *N*1 is substituted by *N*2 in the germline), we expect that *f*_*N*1_∼0 and *f*_*N*2_∼1.
- if a subject is heterozygous by *N*1*/N*2, we expect that *f*_*N*1_∼0.5 and *f*_*N*2_∼0.5.
- if the germline nucleotide *N*1 is replaced by a frequent SHM represented by *N*2 (with frequency at least 50%), we expect that *f*_*N*1_ ≤ *freq*.

We classified a position *P* in a gene *G* as a GSV if *f*_*N*1_ ≤ *freq* for at least one subject.

Each *GSV* (represented by a position *P* in a gene *G*, and nucleotides *N*1 and *N*2) was encoded as (*P, G, N*1*/N*2). Figure 3B illustrates the procedure for identifying GSVs using examples of a GSV (126, IGHV1-7, A/G) that represented a known germline variation, a GSV (167, IGHV1-10, G/A) that represents a likely frequent SHM, as well as a non-GSV (180, IGHV1-27, C/T).

Fractions *f*_*N*1_ for position 126 in IGHV1-7 vary from 0.01 to 0.98 and form a trimodal distribution. Since GSV (126, IGHV1-7, A/G) is a known germline variant, the three modes correspond to the homozygous states AA and GG, and a heterozygous state AG. (126, IGHV1-7, A/G) is classified as a GSV because the minimum of fractions *f*_*N*1_ across all individuals (0.01) does not exceed the default frequency threshold *freq*=0.55. (167, IGHV1-10, G/A) was classified as a GSV because fractions *f*_*N*1_ for position 167 of IGHV1-10 vary from 0.54 to 0.86 (likely frequent SHM). In contrast, (180, IGHV1-27, C/T) was classified as non-GSV since fractions *f*_*N*1_ for position 180 in IGHV1-27 vary from 0.78 to 0.95 and did not fall below the frequency threshold.

In total, we classified 52 GSVs in seven V genes: IGHV1-7 (8 GSVs), IGHV1-10 (10), IGHV1-14 (3), IGHV1-17 (7), IGHV1-20 (8), IGHV1-21 (8), and IGHV1-27 (8). 17 out of 52 GSVs represent known germline variations (Figure S1).

### Generating the genotype matrix

For each GSV (*P, G, N*1/*N*2) in each animal, IgQTL computes the *R-ratio* as *R* = *f*_*N*1_ / (*f*_*N*1_ + *f*_*N*2_). The *R*-ratio represents a more flexible and expressive alternative to the conventional binary description of SNP states (e.g., A/A or A/C) because it enables description of SHMs and their relative abundance. The *R*-ratios also distinguish subjects that are heterozygous by the same pair of alleles but have different expression profiles for these alleles (e.g., 80%-20% vs 50%-50%).

We refer to a 52-mer vector of all *R*-ratios for a given animal (across all GSVs) as its *IGenotype*. We further analyzed IGenotypes across all 204 animals for finding their correlations with various phenotypes. The IGenotypes of 204 animals across 52 GSVs form a 52×204 *IGenotype matrix*, an analog of a genotype matrix that describes both genomic SNPs and SHMs (Figure 4A).

**Figure 4.**
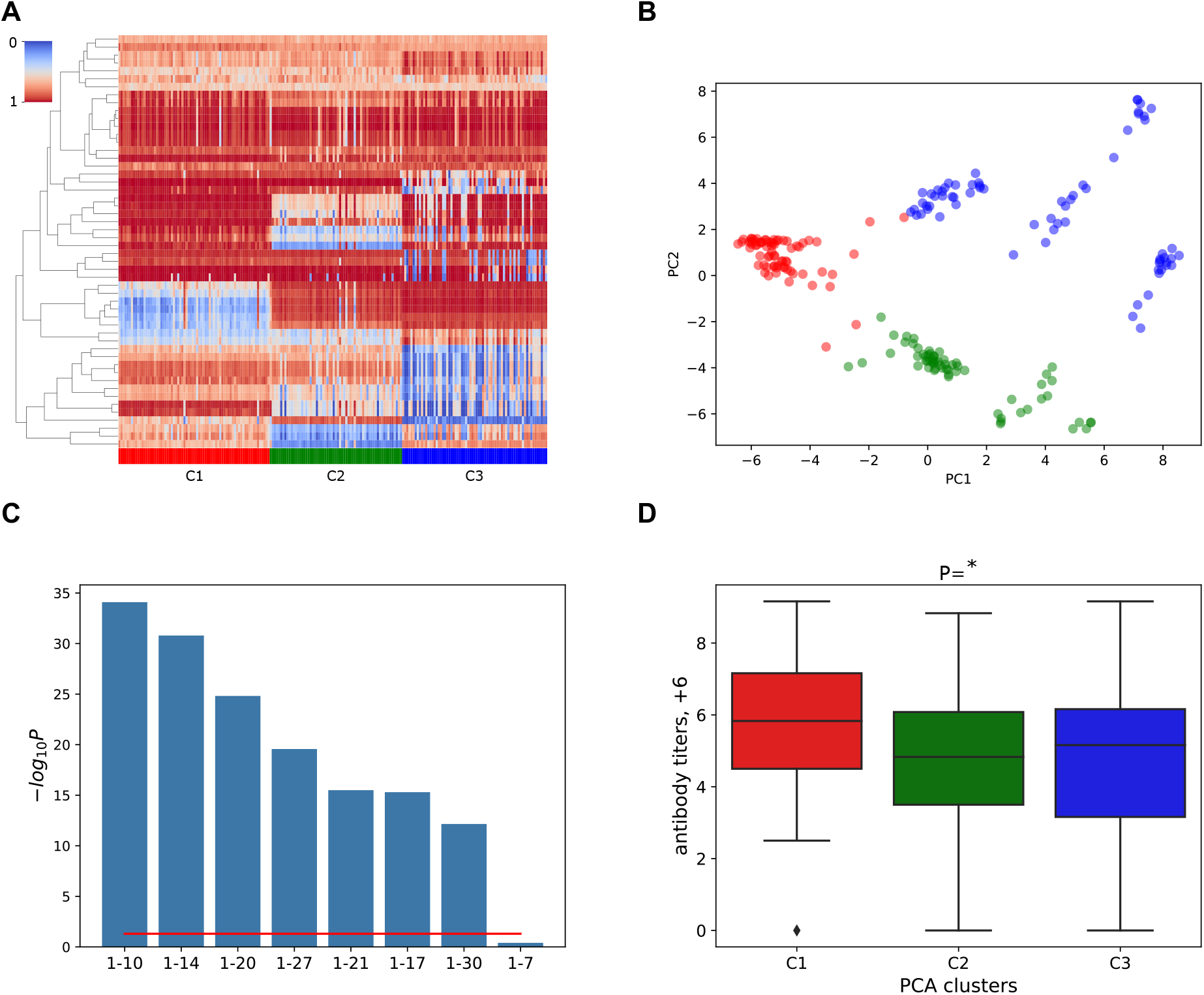
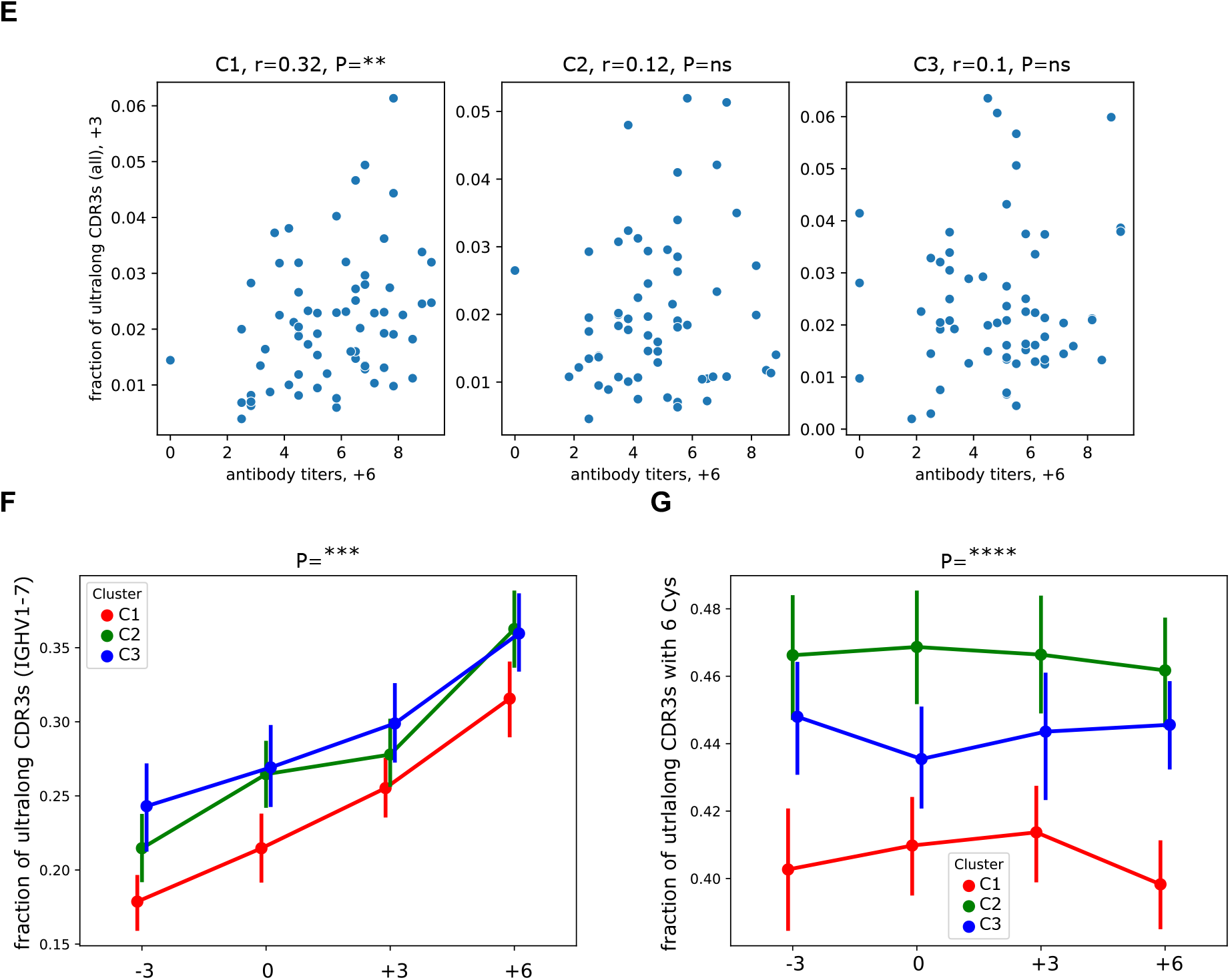
GSVs of V genes are associated with gene usages and antibody titers. (*A*) The 52×204 IGenotype matrix for 52 GSVs of V genes across 204 calves. Rows represent GSVs, columns represent animals. Rows are ordered using the hierarchical clustering, columns are ordered according to the three clusters found using PCA, followed by *k*-means clustering. Three clusters C1, C2, and C3 are shown in red, green, and blue in the lower horizontal panel, respectively. The order of animals within a cluster is chosen arbitrarily. *R*-ratios vary from 0 (blue) to 1 (red). (*B*) Principal components 1 and 2 of the IGenotype matrix shown in (*A*). Three identified clusters are shown in red, green, and blue. (*C*) Likelihoods of association P-values between three PCA clusters and usages of V genes. Usages are computed in the combined datasets. Likelihood is computed as the negative logarithm of the P-value to the base of 10. Genes are shown in the descending order of likelihoods. The red line corresponds to P=0.05. (*D*) Antibody titers at time point “+6” for three PCA clusters. (*E*) Antibody titers at time point “+6” vs fractions of ultralong CDR3s in all CDR3s at time point “+3” across clusters C1–C3. The Pearson correlations (*r*) and P-values (P) are shown at the top of the panel. (*F*) Fractions of ultralong CDR3s among all CDR3s derived from IGHV1-7 in clusters C1, C2, and C3 at four time points. (*G*) Fractions of ultralong CDR3s with six cysteines among all ultralong CDR3s in clusters C1, C2, and C3 at four time points. Vertical lines in (*F*) and (*G*) show 95% confidence intervals.

### Clustering animals with similar IGenotypes

We clustered animals into groups with similar IGenotypes (these groups represent analogs of common genotypes) by applying the principal component analysis (PCA) to the IGenotype matrix. Iterative *k*-means clustering of the first two principal components with *k* from 2 to 10 followed by the elbow method (Thorndike, 1953) reveals that *k*=3 provides the optimal decomposition of 204 animals (Figure 4B, Figure S2). Although decompositions into more clusters resulted in similar values of inertia (Figure S2), we focused our analysis on *k*=3 because it simplified further statistical analysis and allowed us to apply popular statistical methods such as the Kruskal-Wallis test (Kruskal and Wallis, 1952).

We say that the computed clusters are *associated* with the *R*-ratios (or usages/titers/fractions of ultralong CDR3s) if the differences between distributions of *R*-ratios (or usages/titers/fractions of ultralong CDR3s) across these clusters are statistically significant. The computed clusters C1, C2, and C3 are associated with the *R*-ratios for 47 out of 52 GSVs, including 16 out of 17 known germline variations (Figure S3). We thus conclude that the decomposition of 204 calves into clusters C1–C3 is driven by multiple *linked* GSVs (GSVs with correlated *R*-ratios) that represent common genotypes of V genes.

### GSVs explain variance in usages of V genes and antibody titers

Clusters C1–C3 are associated with usages of all highly-used V genes except for IGHV1-7 (Figure 4C, Figure S4). Figure 4D shows that the clusters are also associated with antibody titers collected at time point “+6”: cluster C1 has higher antibody titers compared to clusters C2 and C3 with P=0.017 according to the Kruskal-Wallis test. This observation suggests that IGenotypes of V genes are associated with the response to the BRD vaccination. Antibody titers collected at three other time points do not have statistically significant associations with clusters C1– C3.

Since the fractions of ultralong antibodies are not associated with clusters C1–C3 (Figure S5), we hypothesize that generation of ultralong antibodies is not specific to genotypes described by the revealed clusters but rather is a general feature of cattle antibody repertoires. However, our analysis revealed subtle correlations between genotypes and some features of ultralong CDR3s. Figure 4E shows that clusters C1– C3 partially explain the variance in Figure 2D: the fraction of ultralong CDR3s among all CDR3s at the time point “+3” positively correlates with titers at the time point “+6” only for animals from the cluster C1 (*r*=0.32, P=0.0072). Similar correlations do not exist and are not statistically significant for clusters C2 and C3 (*r*=0.12 and *r*=0.10, respectively). We thus assume that ultralong CDR3s from the cluster C1 work better in response to the BRD vaccine.

Figure 4F and G show that animals from the cluster C1 are characterized by a lower initial fraction of ultralong CDR3s (in all CDR3s derived from IGHV1-7) and a lower fraction of ultralong CDR3s with six cysteines (important for knob formation) as compared to animals from clusters C2 and C3 (P-values for the cluster variable are 3.96×10^−4^ and 2.63×10^−5^, respectively). Since the initial number of cysteines in ultralong CDR3s is four (Wang et al., 2013), a higher number of cysteines suggests that ultralong CDR3s of animals from clusters C2 and C3 underwent more extensive affinity maturation before the BRD vaccination compared to animals from the cluster C1. Since the cluster C1 is associated with higher titers after the second BRD vaccination, we extend the hypothesis about pre-existing immunity and suggest that it might partially consist of mature ultralong CDR3s (with six cysteines) generated before the vaccinations in animals from clusters C2 and C3. However, since titers at time points “–3” and “+6” are anticorrelated in all clusters (Figure S6), we also suggest that mature ultralong antibodies might be not the only component of the pre-existing immunity. Further exploration of cattle antibody repertoires would help to understand the origin of the pre-existing immunity (e.g., maternal antibodies or microbiota) and its impact on the BRD vaccination.

### A GSV at position 148 in IGHV1-7 is associated with the fraction of ultralong CDR3s

The fraction of ultralong CDR3s among all antibodies varies between 0.0033 and 0.0543 in the combined datasets (Figure S7A). The fraction of ultralong CDR3s limited to CDR3s derived from IGHV1-7 varies from 0.09 to 0.64 in the combined datasets (Figure S7B). Since the clusters C1–C3 are not associated with fractions of ultralong CDR3 antibodies (Figure S5), we also examined potential associations of individual GSVs with fractions of ultralong CDR3s.

The GSV (148, IGHV1-7, A/G) has the most significant association with the fraction of ultralong CDR3s whether computed with all antibodies or limited to IGHV1-7 containing antibodies: P=3.03×10^−28^ and P=1.49×10^−47^, respectively (Figure 5A, B). P-values were computed using the linear regression model. The GSV with the next most significant association was GSV (144, IGHV1-17, C/T), but this significance was many orders of magnitude lower (P=0.0016). The closest IGHV1-7 containing GSV in terms of significance was GSV (71, IGHV1-7, C/T) which also has association that is orders of magnitude lower (P=0.0002). Thus, GSV (148, IGHV1-7, A/G) is unique since its association P-values are many orders of magnitude lower than association P-values of all other GSVs.

**Figure 5.**
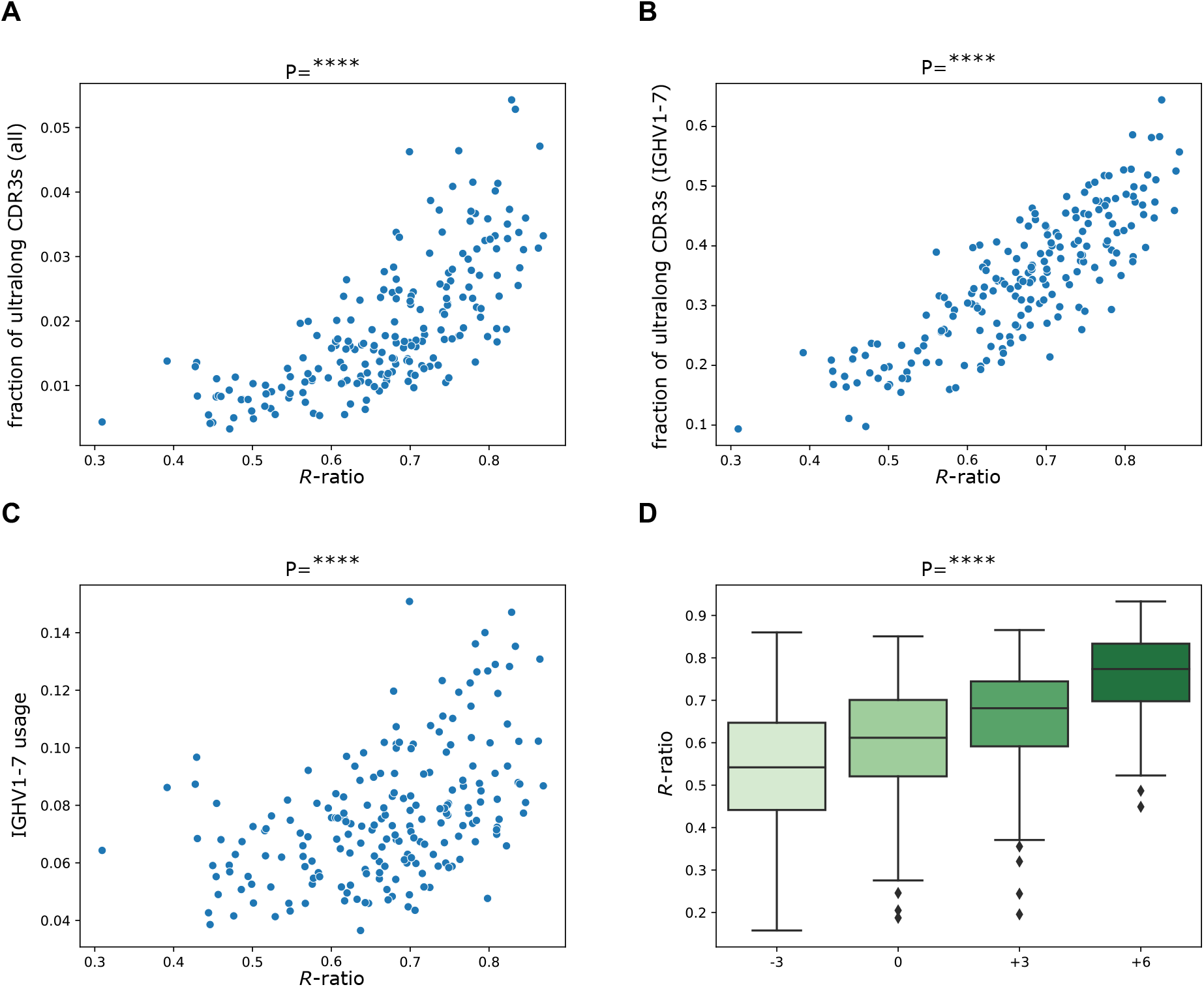
GSV at position 148 of IGHV1-7 is associated with production of ultralong antibodies. (*A*) Fractions of ultralong CDR3s in all CDR3s vs *R*-ratios of the GSV G148A in IGHV1-7. (*B*) Fractions of ultralong CDR3s in all CDR3s derived from IGHV1-7 vs *R*-ratios of the GSV G148A in IGHV1-7. (*C*) Usages of IGHV1-7 in the combined dataset vs *R*-ratios of the GSV G148A at position 148 in IGHV1-7. (*D*) Distributions of *R*-ratios for position 148 in IGHV1-7 at four time points.

This GSV (that we refer to as G148A for brevity) also has the most significant association with the usage of IGHV1-7 in the combined dataset (P=6.11×10^−17^, the linear regression model): the higher is the fraction of nucleotide A at position 148 of IGHV1-7, the higher is the usage of IGHV1-7 (Figure 5C). Figure 5D shows that *R*-ratios of the GSV G148A grow after both vaccinations (P=3.36×10^−204^). This position was not previously identified as a germline variation (Figure 3B), the known alleles of IGHV1-7 have a nucleotide G at position 148 classified as the second popular nucleotide *N*2 by our analysis (Table S2). The *R*-ratios of GSV G148A are not associated with clusters C1–C3, suggesting that the GSV is not linked with 47 GSVs which *R*-ratios are associated by clusters C1–C3 (Figure S3). The *R*-ratios for this GSV varies from 0.31 to 0.87, indicating that both nucleotides A and G are always present in Rep-seq reads in each animal (Figure S9A). Figure S9B also shows that *R*-ratios of the GSV grow similarly for clusters C1–C3. We thus assume that GSV G148A is a frequent SHM that is often selected after vaccinations and is important for generating ultralong antibodies.

### The role of the GSV G148A in ultralong CDR3s

The most abundant nucleotide *N*1=A of the GSV G148A replaces the germline amino acid Gly (encoded by the codon GGT) with the amino acid Ser (encoded by codon AGT) at amino acid position 50. This position represents the last amino acid of the second framework region (FR2) according to the IMGT notation (Lefranc et al., 2003) but is classified as a CDR2 position according to Kabat (Kabat et al, 1979) and Paratome (Kunik et al., 2012) notations. Wang et al., 2013 showed that, unlike Gly at position 50 (referred to as Gly50), Ser at position 50 (referred to as Ser50) can form hydrogen bonds with the conserved Gln at position 97 in the stalk part of an ultralong CDR3. On average, 24.8% and 3.8% of ultralong antibodies derived from IGHV1-7 in the combined datasets contain Ser50 and Gly50, respectively (Figure 6A). In contrast to ultralong antibodies, where Ser50 is six-times more frequent than Gly50, Ser50 and Gly50 appear in similar proportions (30.4% and 24.8%, respectively) in non-ultralong CDR3s derived from IGHV1-7 (Figure 6A).

**Figure 6.**
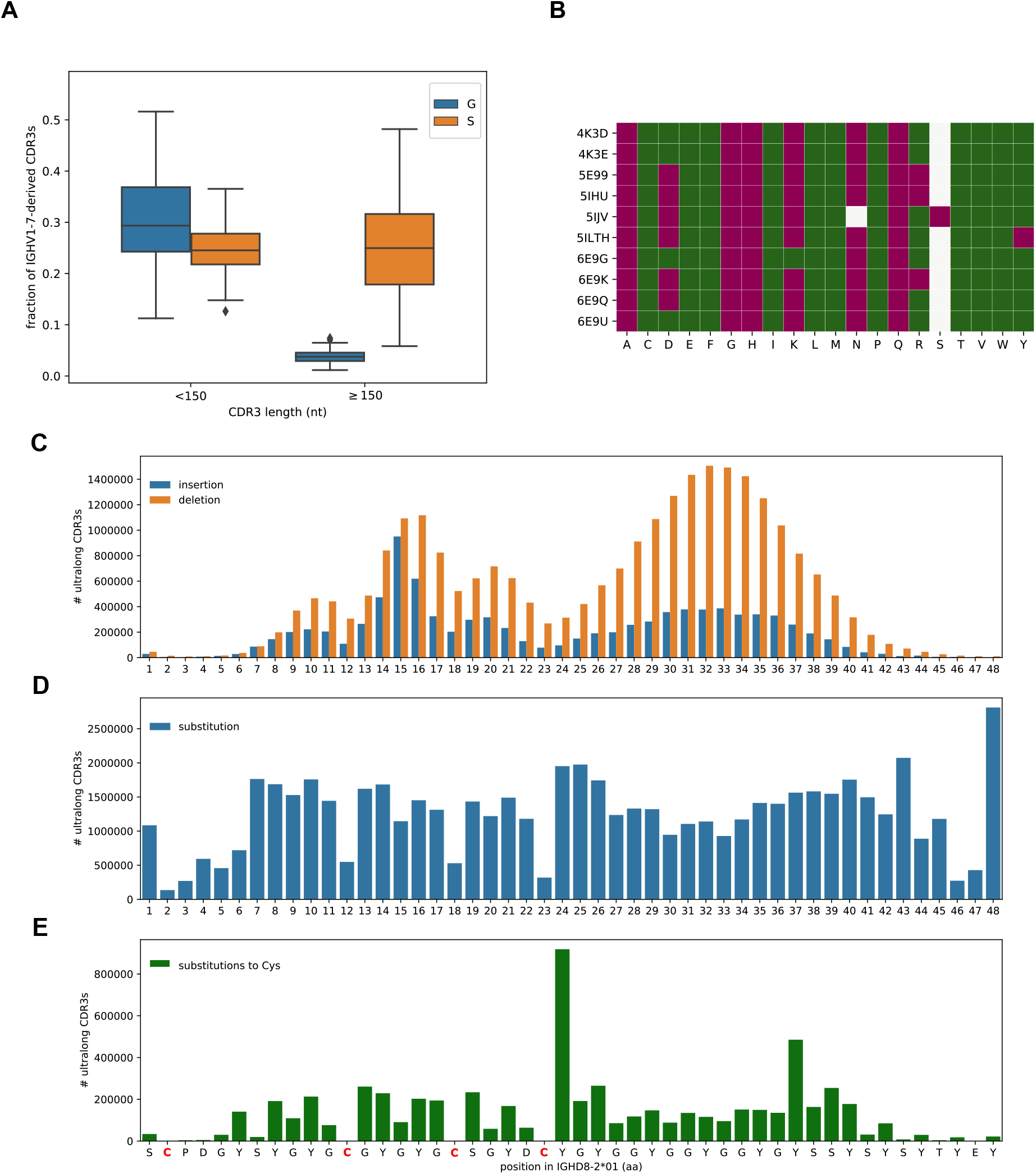
The anatomy of ultralong CDR3s. (*A*) Fractions of non-ultralong CDR3s (lengths below 150 nt) and ultralong CDR3s derived from IGHV1-7 with amino acids Gly (blue) and Ser (orange) at amino acid position 50. Fractions are computed in the combined datasets. (*B*) Impact of substitutions at position 50 in ten crystallized antibodies predicted by the I-Mutant2.0 tool. Accession IDs of antibody structures are shown on the left. White cells show amino acids at position 50 present in structures (N or S). A green (red) cell (*Ab, AA*) indicates that mutation to amino acid *AA* is predicted to increase (decrease) stability of antibody structure *Ab*. (*C*–*E*) IGHD8-2 is a template for generating cysteines through SHMs. The germline sequence of IGHD8-2*01 is shown at the bottom with four cysteines shown in red. (*C*) The bar plot shows the number of ultralong CDR3s that have insertions (blue) and deletions (orange) at a given position of IGHD8-2*01. The position of insertion is defined as the position in the germline sequence that precedes the insertion. The average lengths of insertions and deletions in IGHD8-2 are 1.6 and 2.4 aa, respectively. (*D*) The bar plot shows the number of ultralong CDR3s that have a substitution at a given position of IGHD8-2*01. (*E*) The bar plot shows the number of ultralong CDR3s that have a substitution into cysteine at a given position of IGHD8-2*01.

We also analyzed thirteen 3-D structures of crystallized bovine antibodies (reported by Wang et al., 2013, Stanfield et al., 2016, and Dong et al., 2019) available in the Protein Data Bank (Berman et al., 2000). None of the known ultralong antibodies have the germline Gly at position 50: all but one of them have Ser at position 50 (Figure S10). We further applied the I-Mutant2.0 tool (Capriotti et al., 2005) to analyze the effect of substitutions at this position on the stability of the analyzed antibodies. I-Mutant2.0 generated prediction for only 10 out of 13 analyzed antibodies (three structures were processed with errors) and it turned out that substitution of Ser by the germline amino acid Gly decreases antibody stability for all ten of them (Figure 6B). We thus assume that amino acid Ser at position 50 is critically important for maintaining the structure of ultralong antibodies.

### Surprising features of ultralong CDR3s

All ultralong CDR3s are generated through recombination of a 148 nt long D gene IGHD8-2 that encodes four cysteines in its third open reading frame and contains 39 codons (in the same frame) differing from the cysteine-encoding codons by a single nucleotide, providing multiple opportunities for generating novel disulfide bonds in an ultralong antibody by somatic mutations of non-cysteines into cysteine. During VDJ recombination, all short IGHD genes undergo intensive exonuclease removals that contribute to the overall diversity of an antibody repertoire (Murphy et al., 2016). To understand the recombination properties of the long IGHD8-2 gene, we collected 2,855,428 distinct ultralong CDR3s across all individuals, aligned them to the IGHD8-2*01 gene, and identified positions corresponding to substitutions, insertions, and deletions. In surprising contrast to the short D genes, the first six and the last three amino acid positions of IGHD8-2 do not accumulate insertions and deletions (Figure 6C) and only the first and last positions undergo substantial numbers of substitutions (Figure 6D). Therefore, in contrast to the short D genes, IGHD8-2 does not undergo extensive truncations from both sides.

The distribution of observed indels in the IGHD8-2*01 segment shows an uneven distribution throughout the middle portion, between positions 6 and 45 (Figure 6C). Relatively increased numbers of indels can be detected in the region between positions 14–17 and downstream from position 25 to 40, with deletions much more common than insertions in the latter. We note that the positions prone to indels do not include those encoding four germline cysteines (positions 2, 12, 18, and 23). Furthermore, the germline cysteine codons accumulate ∼5 times fewer substitutions compared to other positions of IGHD8-2 (Figure 6D). Thus, most ultralong CDR3s preserve germline cysteines of IGHD8-2. The introduction of additional cysteines in the ultralong CDR3 by SHM are most commonly the result of substitutions at position 24 (∼4 times more common than the average at other positions; Figure 6E).

## Discussion

### Longitudinal study of cattle antibody repertoires developed in response to the BRD vaccine

We conducted a personalized immunogenomics study of 204 calves to analyze the efficacy of the BRD vaccine. Our analysis showed that the BRD vaccinations increase both the usage of IGHV1-7 and the fraction of ultralong antibodies, suggesting that ultralong antibodies play an important role in immune responses against antigens of the BRD vaccine. It also showed that antibody titers measured after the booster vaccination are weakly correlated with the usage of IGHV1-7 and the fraction of ultralong antibodies. Usages of other cattle IGHV genes are not associated with antibody titers before and after the BRD vaccination. We also showed that antibody titers before the initial vaccination anticorrelate with titers after the booster vaccination. This suggests that pre-existing immunity to BRD may prevent successful development of the immune response to the BRD vaccine.

### The IgQTL analysis of antibody repertoires

Although the analysis of eQTLs in genomic studies is well developed, there are still no tools for analyzing IgQTLs in immunogenomics datasets. We developed an IgQTL tool for detecting important germline and somatic variations or GSVs (based on analyzing Rep-seq data), applied it to identify GSVs in cattle IGHV genes, and found their associations with various phenotypes (gene usages, fractions of ultralong antibodies, and antibody titers). Our analysis demonstrates that IgQTL can be used for analyzing antigen-specific antibody responses in a population. Although it has only been tested on cattle Rep-seq datasets, it can be applied to any vertebrate species, including humans, and thus improve our understanding of the specifics of adaptive immune responses associated with various antigens.

### SHM G148A in IGHV1-7 is important for generating ultralong antibodies

Analysis of the identified GSVs revealed that a GSV G148A in IGHV1-7 is strongly associated with both the usage of IGHV1-7 and the fraction of ultralong antibodies. This GSV results in a substitution of the germline amino acid Gly into Ser that is specific to ultralong antibodies. While non-ultralong antibodies derived from IGHV1-7 have similar fractions of Gly and Ser, the ultralong antibodies have a highly elevated fraction of Ser as compared to the fraction of Gly. Wang et al., 2013 showed that Ser encoded by this GSV forms a hydrogen bond with the conservative Gln at position 95 in the stalk region of an ultralong CDR3. We thus assume that the GSV G148A is not specific to responses induced by the BRD vaccine but rather is a general feature of ultralong antibodies. Different patterns of the GSV G148A in IGHV1-7 in ultralong and non-ultralong antibodies suggest that it represents a frequent SHM rather than a novel germline variation. Further investigation of the origin and the role of this GSV will likely require paired WGS and Rep-seq datasets, as well as analyzing the 3-D structures of ultralong antibodies.

### Germline variations and SHMs explain variance in titers

Further analysis of GSVs revealed three clusters (C1, C2, and C3) representing common genotypes of IGHV genes. The detected clusters are associated with usages of all highly-used IGHV genes except for IGHV1-7. The cluster C1 is associated with higher titers after the booster vaccination and a higher correlation between the final titers and the fraction ultralong CDR3s compared to clusters C2 and C3. The cluster C1 is also characterized by a significantly lower fraction of ultralong CDR3s before the vaccination and a lower fraction of ultralong CDR3s with six cysteines. Since the initial number of cysteines in ultralong CDR3s is four (Wang et al., 2013), a higher number of cysteines indicates that ultralong CDR3s of animals from clusters C2 and C3 underwent more extensive affinity maturation compared to animals from the cluster C1. We conjectured that the pre-existing immunity partially consists of “mature” ultralong CDR3s (with six cysteines) detected before the vaccinations in animals from clusters C2 and C3. However, further exploration of cattle antibody repertoires is needed for understanding the origin of the pre-existing immunity and its impact on the BRD vaccination.

### Further analysis of antibody responses to BRD

The GSVs detected by IgQTL might be useful for identifying “vaccine-ready” animals (e.g., animals from the cluster C1) capable of mounting efficient responses to the BRD vaccine. Further studies examining GSVs and their relationship with developing antibody repertoires in response to vaccination have a potential to identify animals with successful responses to the BRD vaccine and thus contribute to the ongoing selection strategies by including not only genomic but also immunogenomic traits.

Our study has revealed that ultralong antibodies play an important role in the antibody response against BRD antigens. Although our study has already combined experimental (Rep-Seq) and computational approaches, these approaches would further benefit from functional experiments aimed at identification of ultralong antibodies that bind BRD antigens. Such experiments (and further antibody engineering analysis) represent the topic of a follow-up paper.

## Methods

### Sample preparation

The Iowa State University Animal Care and Use Committee (IACUC) approved all animal work before the study was conducted. Purebred American Angus calves (*n* = 204) were vaccinated with a modified live vaccine (Bovi-Shield Gold 5; Zoetis Inc., Parsippany N.J.) containing antigens of 4 viruses associated with bovine respiratory disease as described (Kramer et al., 2017). A second booster vaccination was applied three weeks later. Bovine whole blood was collected and stored in Tempus TM blood RNA tubes (Applied Biosystems, Foster City, CA). Collections occurred at four-time points; three weeks prior to vaccination, at vaccination, three weeks post-vaccination (at booster vaccination), and 3 weeks after booster vaccination. RNA was isolated using the Tempus TM spin RNA isolation kit as recommended by the manufacturer (Applied Biosystems, Foster City, CA). Antibody titers were quantified as previously describe (Kramer et al., 2017).

### Repertoire sequencing

The RNA was converted to cDNA with the Clontech SMARTer kit (Takara Bio USA; Mountain View, CA) using the modified (Switching Mechanism At 5’ End of RNA Transcript) PCR cDNA synthesis protocol (Clontech) and oligonucleotides as described below (Table S3). 2 µg of RNA in 11 µl was mixed with 1 µl of 12 µM SMARTer IIA Oligonucleotide. The mixture was incubated at 72 °C for 30 min and then 42 °C for 2 min. 9 µl of master mix, containing 87747 primer, 5X first-strand buffer, 0.1M DTT, 10 mM dNTP mix, Recombinant RNasin Ribonuclease inhibitor (Promega), and SuperScript II reverse transcriptase (Invitrogen) was added to the mixture and incubated at 42 °C for 90 min and then 70 °C for 10 min. Since primer 87747 (3’ SMARTer CDS IIA) targeted on the 5’end of V H leader sequence, the cDNA synthesis produced high quality double strand IgG heavy chain cDNA.

Illumina amplicon libraries were designed to cover the IgG heavy chain variable region (FR1-4 and CDR1-3) as previously described (Larsen and Smith, 2012). Libraries were constructed by employing two primers that targeted 3’ end of the leader region (87934 primer) and 5’ end of the IgG heavy chain constant (CH1) region (87935 primer). Table S3 presents information about the primers that were used for this study. For PCR amplification for libraries, AccuPrime Tag DNA polymerase High Fidelity (Invitrogen) with initial denaturation at 94 °C for 2 min, 33 cycles of 94 °C for 15 sec, 64 °C for 15 sec, 68 °C for 1 min, and final extension 68 °C for 5 min conditions were used. The libraries were purified by using AMPure XP bead-based purification as recommended by the manufacturer (Beckman Coulter, Brea CA). The concentration of libraries was determined by quantitative PCR using a NEBNext library Quant kit for Illumina (New England BioLabs, Ipswich MA). The size of libraries and amplicon profiles were determined by the Fragment Analyzer System (Agilent Technologies, Santa Clara CA). Paired end (2×300 cycle) sequencing was performed on a Miseq® sequencer using a 600 cycle v3 Reagent Kit (Illumina Inc., San Diego CA) producing 300 base paired end reads.

### Preprocessing Rep-seq data

Each paired-end read was merged into a single sequence using the PairedReadMerger tool (Safonova et al., 2015). For each resulting sequence, the V gene, the J gene, and the CDR3, contributing to this sequence were inferred using the DiversityAnalyzer tool (Shlemov et al., 2017) based on the cattle germline immunoglobulin genes listed in the IMGT database (Lefranc et al., 2009). To simplify the downstream analysis of gene variations, we kept only the first allele of each V gene and ignored its allele variants. The germline and somatic variations were computed using alignments against the closest V genes reported by the DiversityAnalyzer.

### Statistical analysis

Statistical analysis was performed using Python (version 3.8.5). The Kruskal-Wallis test and the Pearson correlation were computed using the Scipy package (version 1.6.0). The linear regression model and the linear mixed effect model were called from the Statmodels package (version 0.12.2).

### Data Access

Sequencing datasets were deposited to NCBI BioProject under accession number PRJNA607961 (<u>reviewer’s link</u>). Scripts and results are available at GitHub: github.com/yana-safonova/IgQTLgithub.com/yana-safonova/IgQTL.

## Acknowledgements

We are grateful to Vaughn Smider for fruitful discussions and thoughtful comments. We also thank Kristen Kuhn, Jacky Carnahan, and William Thompson for technical support. Mention of trade names or commercial products in this publication is solely for the purpose of providing specific information and does not imply recommendation or endorsement by the U.S. Department of Agriculture. USDA is an equal opportunity provider and employer.

