## Supplemental Materials for "Revealing how variations in antibody repertoires correlate with vaccine responses"

The document includes 10 supplemental figures, 3 supplemental figures, and one supplemental note.

### Supplemental Figures

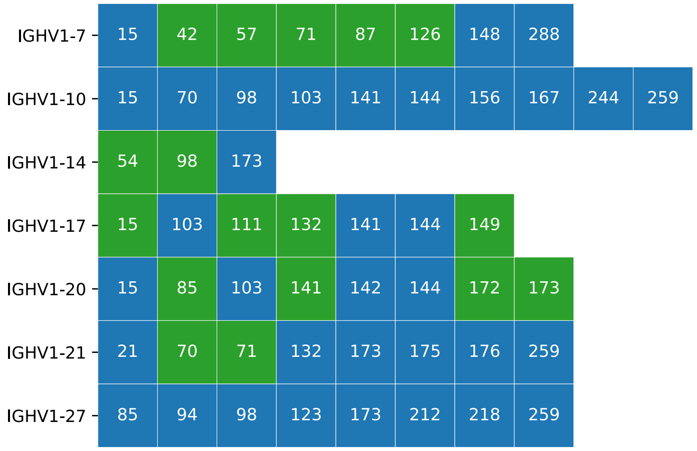


**Figure S1. 52 identified GSVs.** The number within a cell shows the nucleotide position of the GSV in the first IMGT allele of the gene. 17 green cells represent known germline variations. The remaining 35 blue cells represent either still unknown germline variations or frequent SHMs.

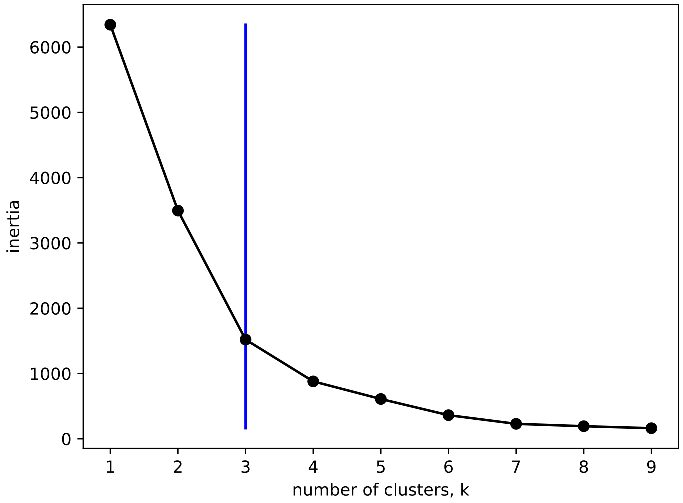


**Figure S2. Inertias of clusters computed via *k*-means clustering for the matrix shown in Figure 4A.** The vertical blue line indicates the optimal number of clusters identified using the elbow method.

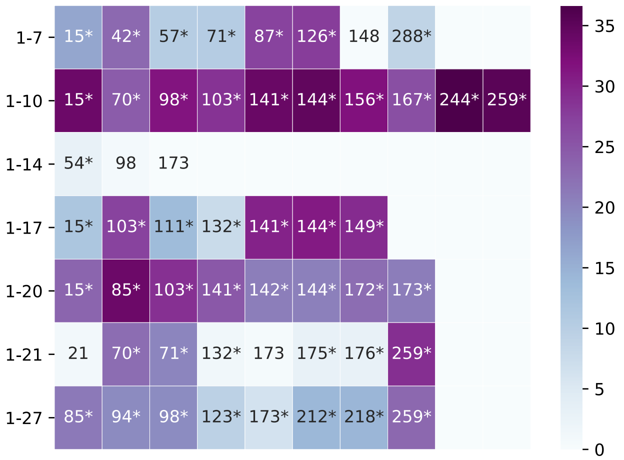


**Figure S3. Associations of *R*-ratios of 52 GSVs of V genes with clusters C1–C3 shown in Figure 4B.** Cells show association likelihoods (= $-\log_{10}P$) varying from 0 (white) to 36 (violet). Variations are labeled by positions in V genes. Statistically significant associations (P<0.05) are additionally labeled with "*".

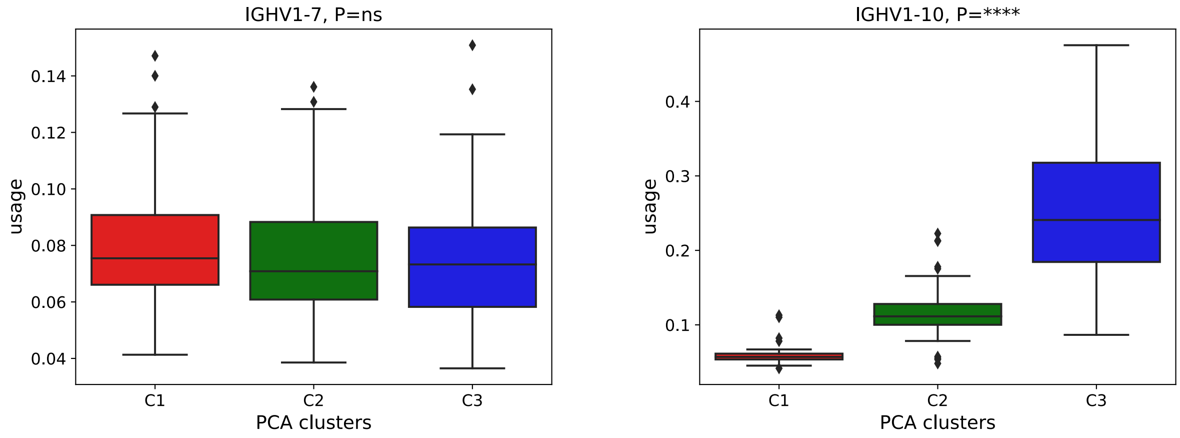

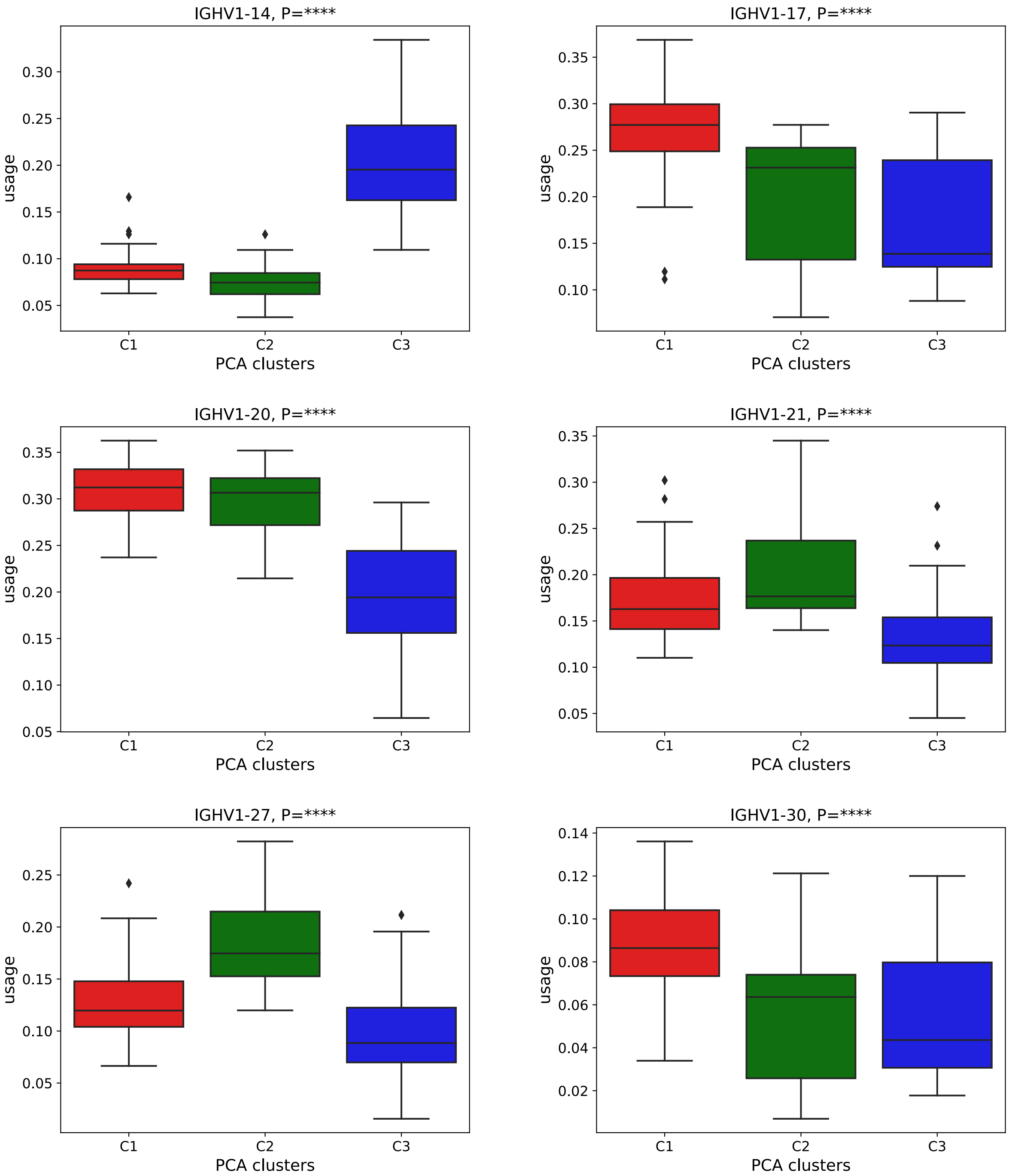


**Figure S4. Distribution of usages of eight V genes in three clusters identified using PCA shown in Figure 4.** P-values were computed using the Kruskal-Wallis test.

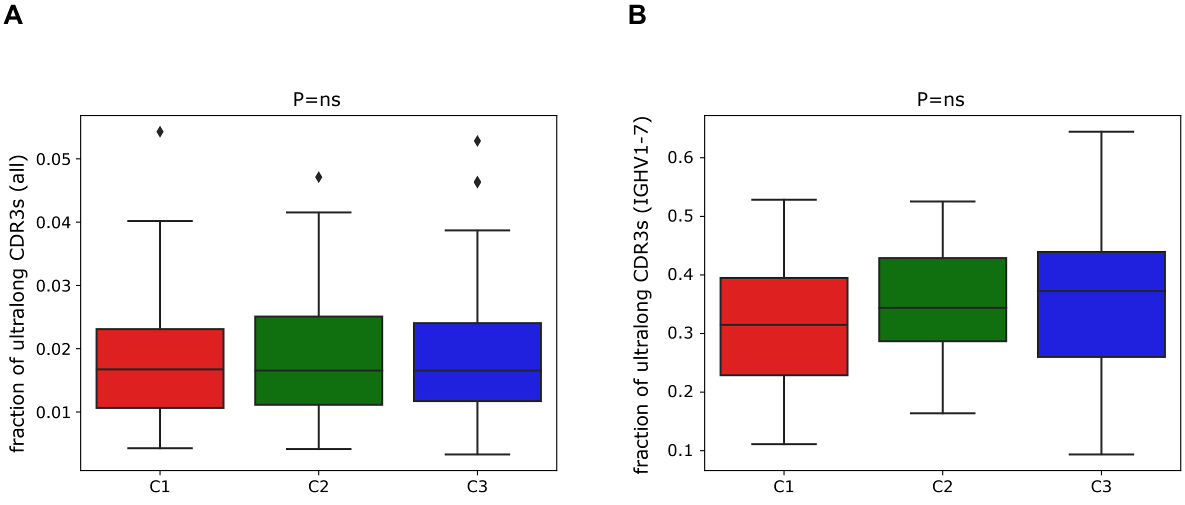


**Figure S5. Fraction of ultralong CDR3s in all CDR3s (*A*) and CDR3s derived from IGHV1-7 only (*B*) across clusters C1, C2, and C3.** P-values were computed using the Kruskal-Wallis test.

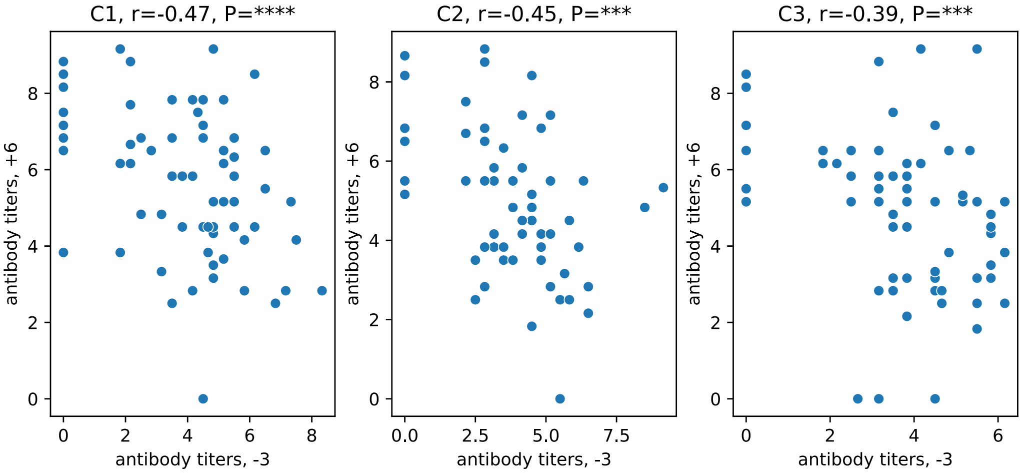


**Figure S6. Antibody titers at time point "–3" vs antibody titers at time point "+6" in clusters C1 (left), C2 (middle), and C3 (right).** The Pearson correlations (*r*) and P-values (P) are shown the top of panels.

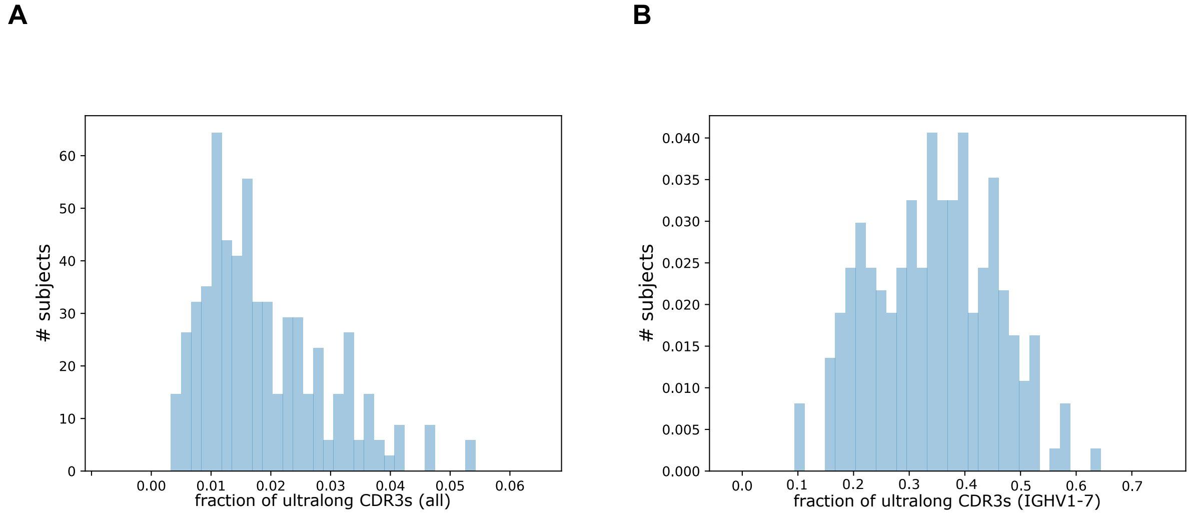


**Figure S7. Fractions of ultralong CDR3s.** (*A*) The distribution of the fractions of ultralong antibodies among all antibodies in the combined datasets. (*B*) The distribution of the fraction of sequences with ultralong antibodies in all antibodies derived from IGHV1-7 in the combined datasets.

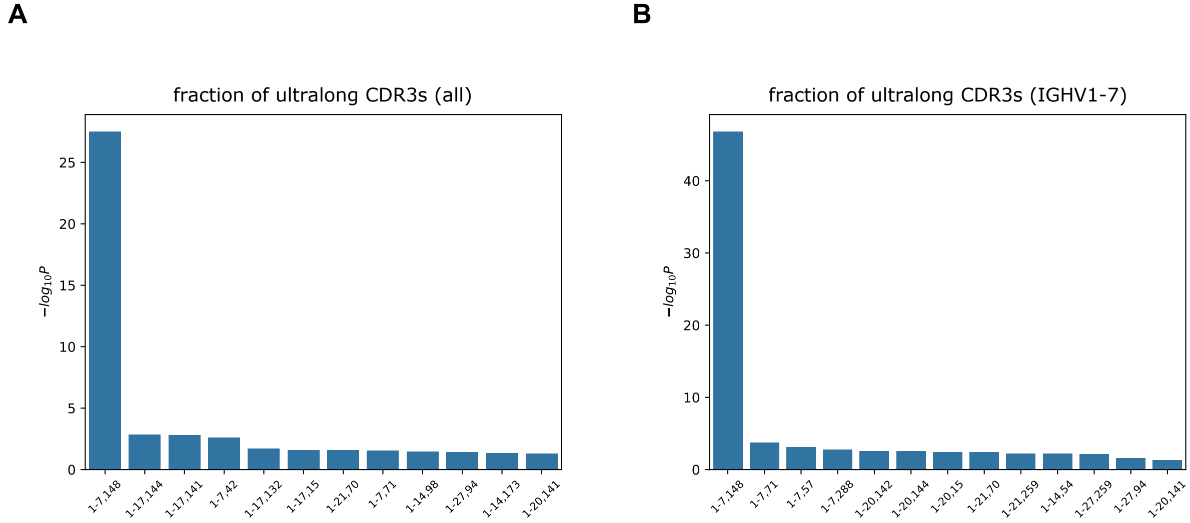


**Figure S8. Likelihoods of association P-values between fractions of ultralong CDR3s and GSVs.** Likelihood is computed as the negative logarithm of the P-value to the base of 10. GSVs are labeled by the V gene and the position in it and shown in the descending order of likelihoods. Only GSVs with P-values below 0.05 are shown. (*A*) and (*B*) show fractions of ultralong CDR3s in all CDR3s and CDR3s derived from IGHV1-7 only, respectively.

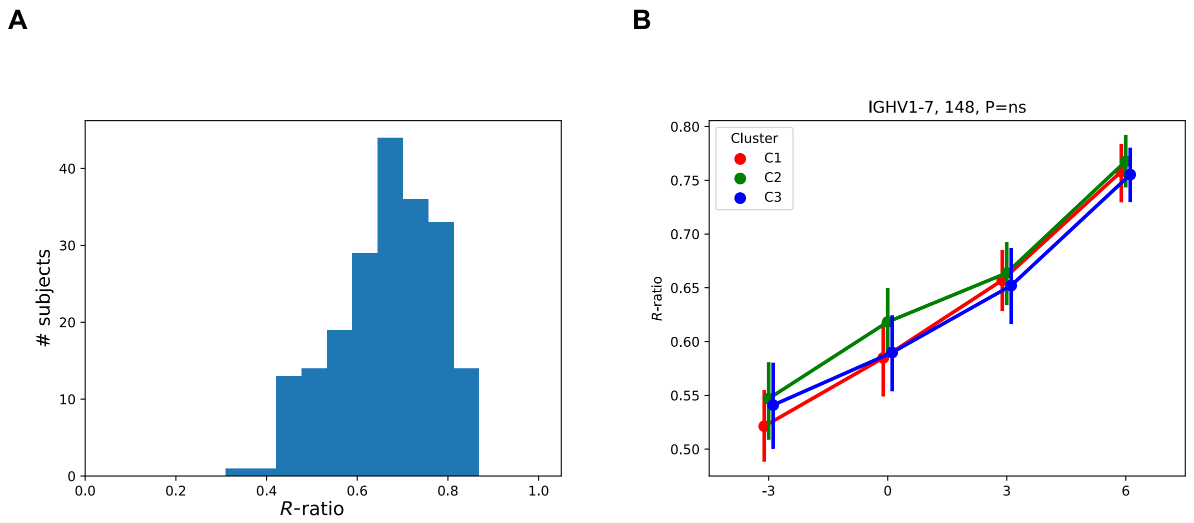


**Figure S9. Characteristics of the GSV (IGHV1-7, 148).** (*A*) The distribution of *R*-ratios at position 148 in IGHV1-7. (*B*) The distribution of *R*-ratios at position 148 across clusters C1–C3. Vertical lines show 95% confidence intervals.

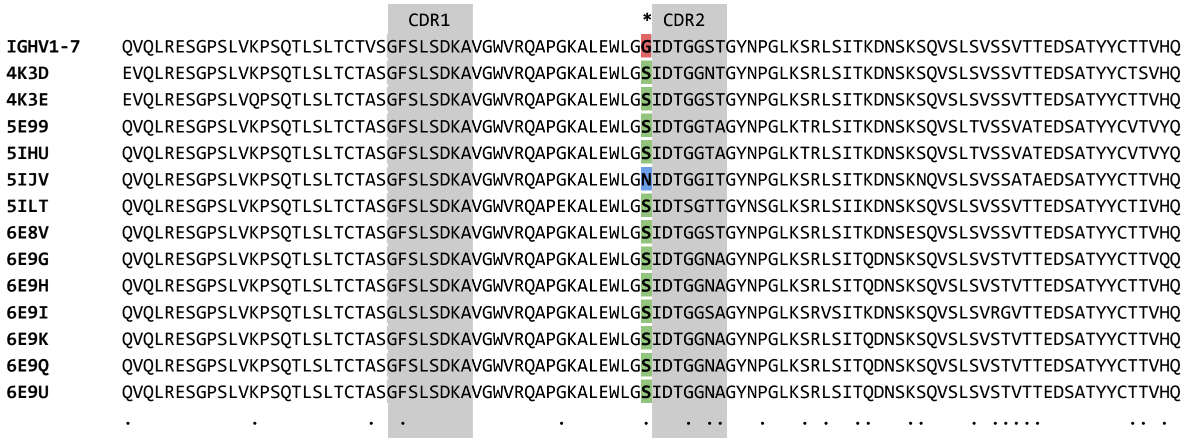


**Figure S10. Known bovine antibodies with ultralong CDR3s support variation at position 148 in IGHV1-7.** Fragments of heavy chain sequences of 13 crystallized bovine antibodies with ultralong CDR3s corresponding to IGHV1-7. Accession numbers of corresponding structures are shown on the left. The germline sequence of IGHV1-7 is shown on the top of the alignment. Amino acid position 50 corresponding to the nucleotide position 148 is marked with "*". Columns with at least one difference from the germline amino acid are marked with "." on the bottom of the alignment. CDR1s and CDR2s (according to the IMGT notation) are shown by grey boxes.

### Supplemental Tables

| Gene | Average usage at "–3" | Average usage at "+6" | Usage fold |
|---|---|---|---|
| IGHV1-7 | 0.06 | 0.11 | 1.83 |
| IGHV1-10 | 0.14 | 0.15 | 1.07 |
| IGHV1-14 | 0.13 | 0.11 | 0.85 |
| IGHV1-17 | 0.22 | 0.21 | 0.95 |
| IGHV1-20 | 0.28 | 0.25 | 0.89 |
| IGHV1-21 | 0.17 | 0.16 | 0.94 |
| IGHV1-27 | 0.13 | 0.14 | 1.08 |
| IGHV1-30 | 0.07 | 0.07 | 1.00 |

**Table S1. Average usages of eight V genes at time points "–3" and "+6".** The column "Usage fold" shows the ratio of the average usages at "+6" to the average usage at "–3" for each V gene.

| Gene | Allele name | Sequence |
|---|---|---|
| IGHV1-7 | 01-IMGT | CAGGTGCAGCTGCGGGAGTCGGGCCCCAGCCTGGTGAAGCCGTCACAGACCCTCTCCCTCACCTGCACGGT<br>CTCTGGATTCTCATTGAGCGACAAGGCTGTAGGCTGGGTCCGCCAGGCTCCAGGGAAGGCGCTGGAGTGGC<br>TCGGTGGTATAGACACTGGTGGAAGCACAGGCTATAACCCAGGCCTGAAATCCCGGCTCAGCATCACCAAG<br>GACAACTCCAAGAGCCAAGTCTCTCTGTCAGTGAGCAGCGTGACAACTGAGGACTCGGCCACATACTACTG<br>TACTACTGTGCACCAGA |
| | 02-IMGT | CAGGTGCAGCTGCGGGAGTCGGGCCCCAGCCTGGTGAAGCCCTCACAGACCCTCTCGCTCACCTGCACGGC<br>CTCTGGATTCTCATTGAGCGACAAGGCTGTAGGCTGGGTCCGCCAGGCTCCAGGGAAGGCGCTGGAGTGGC<br>TCGGTGGTATAGACACTGGTGGAAGCACAGGCTATAACCCAGGCCTGAAATCCCGGCTCAGCATCACCAAG<br>GACAACTCCAAGAGCCAAGTCTCTCTGTCAGTGAGCAGCGTGACAACTGAGGACTCGGCCACATACTACTG<br>TACTACTGTGCACCAGA |
| | Angus | CAGGTGCAGCTGCGGGAGTCGGGCCCCAGCCTGGTGAAGCCGTCACAGACCCTCTCGCTCACCTGCACGGC<br>CTCTGGATTCTCATTAAGCGACAAGGCTGTAGGCTGGGTCCGCCAGGCTCCAGGAAAGGCGCTGGAGTGGC<br>TCGGTGGTATAGACACTGGTGGAAGCACAGGCTATAACCCAGGCCTGAAATCCCGGCTCAGCATCACCAAG<br>GACAACTCCAAGAGCCAAGTCTCTCTGTCAGTGAGCAGCGTGACAACTGAGGACTCGGCCACATACTACTG<br>TACTACTGTGCACCAGA |
| IGHV1-10 | 01-IMGT | CAGGTGCAGCTGCGGGAGTCGGGCCCCAGCCTGGTGAAGCCCTCACAGACCCTCTCCCTCACCTGCACGGT<br>CTCTGGATTCTCATTGAGCAGCTATGGTGTAGGCTGGGTCCGCCAGGCTCCAGGGAAGGCGCTGGAGTGTC<br>TTGGTGGTATAAGTAGTGGTGGAAGCACAGGCTATAACCCAGCCCTGAAATACCGGCTCAGCATCACCAAG<br>GACAACTCCAAGAGCCAAGTCTCTCTGTCACTGAGCAGCGTGACAACTGAGGACACGGCCACATACTACTG<br>TGCGAAGGA |
| | 02-IMGT | CAGGTGCAGCTGCGGGAGTCGGGCCCCAGCCTGGTGAAGCCCTCACAGACCCTCTCCCTCACCTGCACGGT<br>CTCTGGATTCTCATTGAGCAGCTATGGTGTAGGCTGGGTCCGCCAGGCTCCAGGGAAGGCGCTGGAGTGTC<br>TTGGTGGTATAAGTAGTGGTGGAAGCACAGGCTATAACCCAGCCCTGAAATCCCGGCTCAGCATCACCAAG<br>GACAACTCCAAGAGCCAAGTCTCTCTGTCACTGAGCAGCGTGACAACTGAGGACACAGCCACATACTACTG<br>T |
| | Hereford | CAGGTGCAGCTGCGGGAGTCGGGCCCCAGCCTGGTGAAGCCCTCACAGACCCTCTCCCTCACCTGCACGGT<br>CTCTGGATTCTCATTGAGCAGCTATGGTGTAGGCTGGGTCCGCCAGGCTCCAGGGAAGGCGCTGGAGTGTC<br>TTGGTGGTATAAGTAGTGGTGGAAGCACAGGCTATAACCCAGCCCTGAAATCCCGGCTCAGCATCACCAAG |

| | | |
|---|---|---|
| | | GACAACTCCAAGAGCCAAGTCTCTCTGTCACTGAGCAGCGTGACAACTGAGGACACGGCCACATACTACTG<br>TGCGAAGGA |
| IGHV1-14 | 01-IMGT | CAGGTGCAGCTGCGGGAGTCGGGCCCCAGCCTGGTGAAGCCCTCACAGACCCTGTCCCTCACCTGCACGGT<br>CTCTGGATTCTCATTAAGCGATAATAGTGTAGGCTGGGTCCGCCAGGCTCCAGGAAAGGCGCTGGAGTGGC<br>TCGGTGTCATATATAGTGGTGGAAGCACAGGCTATAACCCAGCCCTGAAATCCCGGCTCAGCATCACCAAG<br>GACAACTCCAAGAGCCAAGTCTCTCTATCACTGAGCAGCGTGACAACTGAGGACACGGCCACATACTACTG<br>TGCAAGAGA |
| | 02-IMGT | CAGGTGCAGCTGCGGGAGTCGGGCCCCAGCCTGGTGAAGCCCTCACAGACCCTCTCCCTCACCTGCACGGT<br>CTCTGGATTCTCATTAAGCGATAATAGTGTAGGCTGGGTCCGCCAGGCTCCAGGAAAGGCGCTGGAGTGGC<br>TCGGTGTCATATATAGTGGTGGAAGCACAGGCTATAACCCAGCCCTGAAATCCCGGCTCAGCATCACCAAG<br>GACAACTCCAAGAGCCAAGTCTCTCTATCACTGAGCAGCGTGACAACTGAGGACACGGCCACATACTACTG<br>TGCAAGAGA |
| | 03-IMGT | CAGGTGCAGCTGCGGGAGTCGGGCCCCAGCCTGGTGAAGCCCTCACAGACCCTCTCCCTCACCTGCACGGT<br>CTCTGGATTCTCATTGAGCAGCTATGCTGTAAGCTGGGTCCGCCAGGCTCCAGGGAAGGCGCTGGAGTGGC<br>TCGGTGTCATATATAGTGGTGGAAACACAGGCTATAACCCAGCCCTGAAATCCCGGCTCAGCATCACCAAG<br>GACAACTCCAAGAGCCAAGTCTCTCTATCACTGAGCAGCGTGACAACTGAGGACACAGCCACATACTACTG<br>T |
| IGHV1-17 | 01-IMGT | CAGGTGCAGCTGCGCGAGTCGGGCCCCAGCCTGGTGAAGCCCTCACAGACCCTCTCCCTCACCTGCACGGT<br>CTCTGGATTCTCATTGAGCAGCTATGCTGTAAGCTGGGTCCGCCAGGCTCCAGGGAAGGCTCTGGAGTGGC<br>TTGGTGATATAAGCAGTGGTGGAAGCACAGGCTATAACCCAGCCCTGAAATCCCGGCTCAGCATCACCAAG<br>GACAACTCCAAGAGCCAAGTCTCTCTGTCAGTGAGCAGCGTGACACCTGAGGACACGGCCACATACTACTG<br>TGCGAAGGA |
| | Angus | CAGGTGCAGCTGCGGGAGTCGGGCCCCAGCCTGGTGAAGCCCTCACAGACCCTCTCCCTCACCTGCACGGT<br>CTCTGGATTCTCATTGAGCAGCTATGCTGTAAGCTGGGTCCGCCAGGCTCCAGGGAAGGCGCTGGAGTGGC<br>TTGGTGGTATAAGCAGTGGTGGAAGCACATGCCTATAACCCAGCCCTGAAATCCCGGCTCAGCATCACCAA<br>GGACAACTCCAAGAGCCAAGTCTCTCTGTCAGTGAGCAGCGTGACACCTGAGGACACGGCCACATACTACT<br>GTGCAAAGGA |
| | Hereford | CAGGTGCAGCTGCGCGAGTCGGGCCCCAGCCTGGTGAAGCCCTCACAGACCCTCTCCCTCACCTGCACGGT<br>CTCTGGATTCTCATTGAGCAGCTATGCTGTAAGCTGGGTACGCCAGGCTCCAGGGAAGGCGCTGGAGTGGC<br>TTGGTGATATAAGCAGTGGTGGAAGCACAGGCTATAACCCAGCCCTGAAATCCCGGCTCAGCATCACCAAG<br>GACAACTCCAAGAGCCAAGTCTCTCTGTCAGTGAGCAGCGTGACACCTGAGGACACGGCCACATACTACTG<br>TGCGAAGGA |
| IGHV1-20 | 01-IMGT | CAGGTGCAGCTGCGGGAGTCGGGCCCCAGCCTGGTGAAGCCCTCACAGACCCTCTCCCTCACCTGCACGGT<br>CTCTGGATTCTCACTGAGCAGCTATGCTGTAGGCTGGGTCCGCCAGGCTCCAGGGAAGGCGCTGGAGTGGC<br>TCGGTGGTATAAGCAGTGGTGGAAGCACATACTATAACCCAGCCCTGAAATCCCGGCTCAGCATCACCAAG<br>GACAACTCCAAGAGCCAAGTCTCTCTGTCAGTGAGCAGCGTGACACCTGAGGACACGGCCACATACTACTG<br>TGCGAAGGA |
| | 02-IMGT | CAGGTGCAGCTGCGGGAGTCGGGCCCCAGCCTGGTGAAGCCCTCACAGACCCTCTCCCTCACCTGCACGGT<br>CTCTGGATTCTCATTAAGCGATAATAGTGTAGGCTGGGTCCGCCAGGCTCCAGGGAAGGCGCTGGAGTGCC<br>TCGGTGGTATAAGCAGTGGTGGAAGCACAGGCTATAACCCAGCCCTGAAATCCCGGCTCAGCATCACCAAG<br>GACAACTCCAAGAGCCAAGTCTCTCTGTCAGTGAGCAGCGTGACACCTGAGGACACAGCCACATACTACTG<br>T |
| IGHV1-21 | 01-IMGT | CAGGTGCAGCTGCGGGAGTCGGGCCCCAGCCTGGTGAAGCCCTCACAGACCCTCTCCCTCACCTGCACGAT<br>CTCTGGATTCTCATTGAGCAGCTATGCTGTAGGCTGGGTCCGCCAGGCTCCGGGGAAGGCGCTGGAGTGGG<br>TTGGTGGTATAAGTAGTGGTGGAAGCACATGCCTTAACCCAGCCCTGAAATCCCGGCTCAGCATCACCAAG<br>GACAACTCCAAGAGCCAAGTCTCTCTGTCAGTGAGCAGCGTGACAACTGAGGACACGGCCACATACTACTG<br>TGCGAAGGA |

| | | |
|---|---|---|
| | 02-IMGT | CAGGTGCAGCTGCGGGAGTCGGGCCCCAGCCTGGTGAAGCCCTCACAGACCCTCTCCCTCACCTGCACGATCTCTGGATTCTCATTGAGCAGCTATGCTGTAGGCTGGGTCCGCCAGGCTCCGGGGAAGGCGCTGGAGTGGGTTGGTGGTATAAGTAGTGGTGGAAGCACATGCCTTAACCCAGCCCTGAAATCCCGGCTCAGCATCACCAAGGACAACTCCAAGAGCCAAGTCTCTCTGTCAGTGAGCAGCGTGACAACTGAGGACACAGCCACATACTACTGT |
| | 03-IMGT | AAGGTGCAGCTGCAGGAGTCGGGTCCCAGCCTGGTGAAGCCCTCACAGACCCTCTCCCTCACCTGCACGGCCTCTGGATTCTCATTGAGCAGCTATGCTGTAGGCTGGGTCCGCCAGGCTCCGGGGAAGGCGCTGGAGTGGGTTGGTGGTATAAGTAGTGGTGGAAGCACATGCCTTAACCCAGCCCTGAAATCCCGGCTCAGCATCACCAAGGACAACTCCAAGAGCCAAGTCTCTCTGTCAGTGAGCAGCGTGACAACTGAGGACACAGCCACATACTACTGT |

**Table S2. Known alleles of cattle V genes**. We included V genes that have (i) two and more variants and (ii) average usage exceeding 0.001. V gene sequences were extracted from the IMGT database and genome assemblies of Angus and Hereford breeds available under accession numbers GCF_003369695.1 and CM008188.2, respectively.

| **Primer name** | **Sequence** |
|---|---|
| SMARTer IIA Oligonucleotide | AAGCAGTGGTATCAACGCAGAGTACTTTTTTTTTTTTTTTTTTTTTTTTTTTTVN |
| 87747 | AAGCAGTGGTATCAACGCAGAGTACGCTCSAAGATGAACCCACTGTG |
| 87935 | AATGATACGGCGACCACCGAGATCTACACTCTTTCCCTACACGACGCTCTTCCGATCTTTTCGGGGCTGTGGTGGASG |
| 87934_1 | CAAGCAGAAGACGGCATACGAGATGTCAAAGTGACTGGAGTTCAGACGTGTGCTCTTCCGATCCCCTCCTCTTTGTGCTSTCA |
| 87934_2 | CAAGCAGAAGACGGCATACGAGATGTGACAGTGACTGGAGTTCAGACGTGTGCTCTTCCGATCCCCTCCTCTTTGTGCTSTCA |
| 87934_3 | CAAGCAGAAGACGGCATACGAGATCATCATGTGACTGGAGTTCAGACGTGTGCTCTTCCGATCCCCTCCTCTTTGTGCTSTCA |
| 87934_4 | CAAGCAGAAGACGGCATACGAGATGCCGTTGTGACTGGAGTTCAGACGTGTGCTCTTCCGATCCCCTCCTCTTTGTGCTSTCA |
| 87934_5 | CAAGCAGAAGACGGCATACGAGATCGGCTTGTGACTGGAGTTCAGACGTGTGCTCTTCCGATCCCCTCCTCTTTGTGCTSTCA |
| 87934_6 | CAAGCAGAAGACGGCATACGAGATGTTCCGGTGACTGGAGTTCAGACGTGTGCTCTTCCGATCCCCTCCTCTTTGTGCTSTCA |
| 87934_7 | CAAGCAGAAGACGGCATACGAGATGTAGATGTGACTGGAGTTCAGACGTGTGCTCTTCCGATCCCCTCCTCTTTGTGCTSTCA |
| 87934_8 | CAAGCAGAAGACGGCATACGAGATCCTTCTGTGACTGGAGTTCAGACGTGTGCTCTTCCGATCCCCTCCTCTTTGTGCTSTCA |
| 87934_9 | CAAGCAGAAGACGGCATACGAGATTATGCTGTGACTGGAGTTCAGACGTGTGCTCTTCCGATCCCCTCCTCTTTGTGCTSTCA |

| | |
|---|---|
| 87934_10 | CAAGCAGAAGACGGCATACGAGATCAGGTAGTGACTGGAGTTCAGACGTGTGCTCTTCCGATCCCCTCCTCTTTGTGCTSTCA |
| 87934_11 | CAAGCAGAAGACGGCATACGAGATGTAGACGTGACTGGAGTTCAGACGTGTGCTCTTCCGATCCCCTCCTCTTTGTGCTSTCA |
| 87934_12 | CAAGCAGAAGACGGCATACGAGATGACGTTGTGACTGGAGTTCAGACGTGTGCTCTTCCGATCCCCTCCTCTTTGTGCTSTCA |
| 87934_13 | CAAGCAGAAGACGGCATACGAGATAGCATGGTGACTGGAGTTCAGACGTGTGCTCTTCCGATCCCCTCCTCTTTGTGCTSTCA |
| 87934_14 | CAAGCAGAAGACGGCATACGAGATATTCGAGTGACTGGAGTTCAGACGTGTGCTCTTCCGATCCCCTCCTCTTTGTGCTSTCA |
| 87934_15 | CAAGCAGAAGACGGCATACGAGATCATCGTGTGACTGGAGTTCAGACGTGTGCTCTTCCGATCCCCTCCTCTTTGTGCTSTCA |
| 87934_16 | CAAGCAGAAGACGGCATACGAGATGATCGTGTGACTGGAGTTCAGACGTGTGCTCTTCCGATCCCCTCCTCTTTGTGCTSTCA |

**Table S3. Sequencing primers**. The red nucleotides correspond to oligo (dT) sequences in the primer "SMARTer IIA Oligonucleotide".

### Supplemental Notes

#### Supplemental Note: The relations between pre- and post-vaccination immunity to the BRD vaccine in calves

The simplest vaccination scenario suggests the immunity does not exist before the vaccination and is gained after the vaccination. It results in low and high antibody titers before and after the vaccination, respectively. A more complex case might account for the impact of pre-existing immunity. If a subject has *optimal* pre-existing immunity (immunity that successfully responds to the vaccine), we except that antibody titers before and after the vaccination would be high. However, the BRD vaccination in 204 calves analyzed in this study represents an even more complex case. While the distributions of titers before and after the vaccination have similar mean values (Figure 2A), they anticorrelate: the higher titer before the vaccination, the lower titer after the vaccination, and vice versa (Figure 2B). This cannot be explained by optimal pre-existing immunity because titers before and after vaccinations would be correlated in that case. Thus, we suggest that pre-existing immunity is *suboptimal* and prevents gaining immunity from the vaccine in some calves.
